# LRP-1 promotes tumor progression of triple negative breast cancers by coordinating extracellular matrix remodeling and immune cell infiltration

**DOI:** 10.64898/2026.06.17.732906

**Authors:** Maxence Mocquery-Corre, Lucille Cartier, Abdel-Ilah Aziz, Alexandre Berquand, Julie Clachet, Chloé Jean, Anne-Aurélie Raymond, Hassan El Btaouri, Jean-William Dupuy, Cathy Hachet, Lise Chazée, Katia Savary, Abdelmnim Radoua, Célia Maquin, Eva Brabencova, Camille Boulagnon Rombi, Muriel Barberi-Heyob, Yacine Merrouche, Stéphane Potteaux, Olivier Micheau, Stéphane Dedieu, Jérôme Devy, Jessica Thevenard-Devy

## Abstract

**Background:** Triple-negative breast cancer (TNBC) represents a major clinical challenge due to its aggressiveness, heterogeneity and limited availability of effective targeted therapy. We investigated whether LRP-1, a multifunctional cell-surface endocytic and signaling receptor, contributes to TNBC progression.

**Methods:** Using CRISPR–Cas9, LRP-1-deficient murine 4T1 and human HS578-T TNBC cells were used. Functional consequences were assessed through migration, invasion, and 3D spheroid assays, imaging of focal adhesions and actin organization, atomic force microscopy, and plasmin activity assays. Global molecular reprogramming was analyzed by label-free quantitative proteomics and secretomics. LRP-1-deficient or proficient 4T1 cells were implanted orthotopically in immunocompetent mice; tumor progression was monitored longitudinally while peritumoral collagen architecture and immune microenvironment composition were characterized by second harmonic generation imaging and immunohistochemistry.

**Results:** We show that LRP-1 loss reduces TNBC aggressiveness, as reflected by decreased migration and invasive capacity, reduced spheroid evasion, and significant morphological changes in focal adhesion and actin structure. LRP-1-deficient cells became stiffer and showed lower LOXL-4 levels, while pericellular proteolytic activity remained unchanged, suggesting other proteases mechanism. Multi-omic analysis revealed alterations in extracellular matrix (ECM), epithelial–mesenchymal transition, and inflammatory pathways. *In vivo*, LRP-1-deficiency reduced tumor progression and peritumoral collagen deposition, while increasing CD8^+^ T and Natural Killer cell infiltration, together with a cytokine profiling compatible with a more immune-permissive microenvironment.

**Conclusions:** LRP-1 act as a key contributor in TNBC progression through matrix remodeling, mechano-adaptation, and immune exclusion. Positioning it as a candidate biomarker for TNBC patients who are likely to benefit from stroma-targeting therapies.

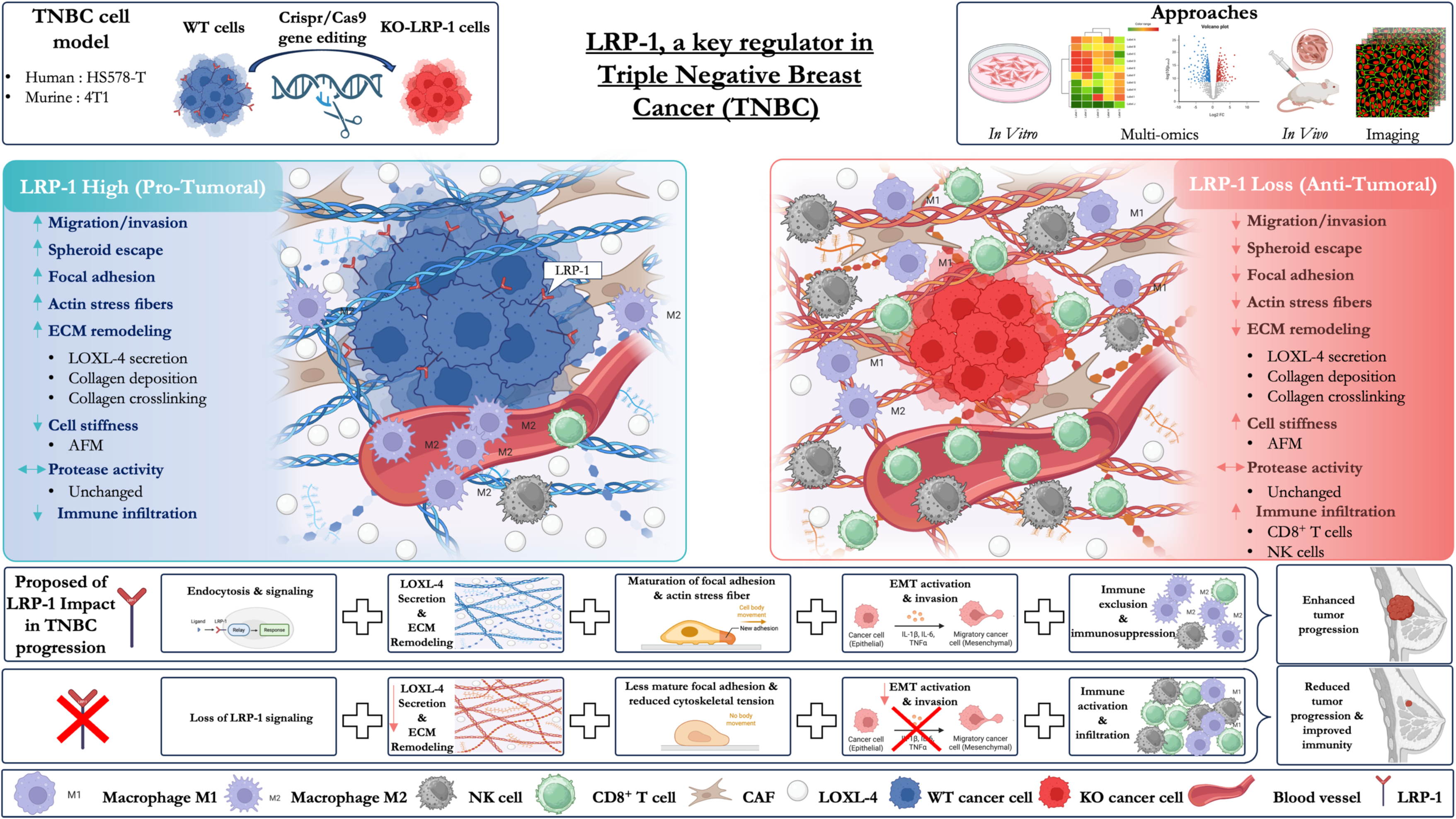

## Introduction

Triple-negative breast cancers (TNBCs), which are distinguished by the low expression of Human Epidermal growth factor receptor 2 (HER2), progesterone, and estrogen receptors, account for roughly 10-15% of all breast cancers[1,2]. They disproportionately affect younger and particular ethnic populations and are characterized by aggressive behavior and high histologic grade[1]. Because TNBC have limited effective targeted therapy, cytotoxic chemotherapies (anthracyclines, taxanes, etc.) remains the clinical standard of care[1], even though new therapies based on immune checkpoint inhibitors (ICIs), anti-PARP agents, and antibody-drug conjugates (ADCs) have emerged in recent years. As a result, TNBC patients typically have worse outcomes and a higher rate of early recurrence than patients with other subtypes[1–3]. Large-scale sequencing studies that have revealed substantial heterogeneity within TNBC, including multiple basal-like and mesenchymal subtypes, have brought attention to the genomic complexity of the disease[1,4]. Despite these advances, there are still few exploitable targets, and novel therapeutic strategies remain limited.

The TNBC tumor microenvironment (TME) is characterized by higher immunogenicity than luminal cancers, as indicated by high Tumor Infiltrating Lymphocytes (TILs) and Programmed death-ligand 1 (PD-L1) [5–7]. Certain TNBC subsets, including germline BRCA1/2 and PD-L1+, respond to immune checkpoint inhibitors or Poly(ADP-ribose)-polymerase (PARP) inhibitors. The tumoral microenvironment (TME), which consists of macrophages, Cancer-associated fibroblast (CAFs), and other immune cells, can either stimulate or inhibit tumor growth [5,8]. Through collagen deposition and Lysyl oxidases (LOX)-mediated crosslinking, extracellular matrix (ECM) remodeling stiffens the matrix that supports metastasis[9–12]. Stiff and aligned fibers enhance directional motility and may also limit immune infiltration[9,13]. Therefore, the interaction between the immune components of the TME and the ECM structure is a crucial factor in the development of TNBC.

Low-density lipoprotein receptor-related protein 1 (LRP-1/CD91) is a large endocytic LDL-receptor family member that is expressed by a variety of cell types[14,15]. 515 kDa α and 85 kDa β subunits are produced from a roughly 600 kDa precursor [14,15]. It binds a range of ligands, such as α2-macroglobulin, ApoE-lipoproteins, and proteases, to mediate endocytosis and clearance [14]. LRP-1 modulates TGF-β, Akt, NF-κB, and MAPK signaling [15,16]; thereby impacting migration and survival in addition to controlling ECM homeostasis and TME inflammation[17]. Consistent with its broad physiological roles, global LRP-1 deletion is embryonically lethal [15].

In many diseases, LRP-1 has context-dependent and often contradictory functions. Through TGF-β/protease regulation, it can cause fibrotic ECM remodeling[18], but it also favors lipoprotein/remnant handling in atheroprotection[19,20] and encourages Aβ clearance in Alzheimer’s disease [21,22]. Similar dual roles in cancer are played by LRP-1, which stimulates the growth of tumors in pancreatic cancer [23], ovarian cancer [24], and glioblastoma [25]. It might, however, offer protection in other situations, like hepatocellular carcinoma [26]. In contrast, its function in melanoma might be influenced by the grade of the cancer or even the other way around[27,28].

Although our team was the first to report a potential pro-tumoral role of LRP-1, its contribution to the different stages of TNBC progression and its distribution across TNBC molecular subtypes remain largely unexplored [16]. In particular, it remains unclear whether LRP-1 is associated with specific TNBC cellular states, how it contributes to the invasive phenotype of TNBC cells, and whether its functions extend beyond tumor-cell intrinsic programs to regulate extracellular matrix (ECM) organization and immune composition within the tumor microenvironment. Addressing these questions is of particular importance given the growing recognition that ECM remodeling, tissue mechanics, and immune exclusion are tightly interconnected processes governing TNBC progression, metastatic dissemination, and therapeutic resistance. Here, through the integration of clinical cohorts, single-cell and spatial transcriptomic datasets, CRISPR/Cas9-mediated gene editing, proteomic and secretomic profiling, biomechanical analyses, and orthotopic syngeneic models, we identify LRP-1 as a key determinant of a mesenchymal, ECM-remodeling TNBC phenotype. We show that LRP-1 promotes tumor invasiveness while coordinating ECM remodeling, collagen deposition, and the establishment of an immune-restrictive microenvironment. Furthermore, LRP-1 depletion is associated with profound alterations in matrix-associated and inflammatory programs, reduced peritumoral collagen accumulation, and enhanced infiltration of cytotoxic immune cells. Collectively, our findings position LRP-1 as a critical regulator of tumor-stroma-immune crosstalk in TNBC and highlight its potential as a therapeutic vulnerability in aggressive breast cancers.

## Materials and methods

Detailed methods are described in Additional_Materiels_1.

## Results

### LRP-1 is heterogeneously expressed in TNBC and selectively enriched within mesenchymal and ECM-remodeling tumor niches

To investigate the clinical and transcriptional landscape of LRP-1 in breast cancer, we first evaluated its protein expression in primary human tumor specimens. Immunohistochemical (IHC) analysis of a clinical cohort of high-grade (SBR grade 3) triple-negative breast cancer tissue sections demonstrated positive, yet highly heterogeneous, LRP-1 protein expression (Fig. 1A). Automated 3,3’-Diaminobenzidine (DAB) quantification revealed substantial inter-tumoral variability in LRP-1 accumulation across varying aggressive histologies, including poorly differentiated invasive ductal carcinomas and metaplastic carcinomas exhibiting sarcomatoid differentiation (Fig. 1B).

**Figure 1:**
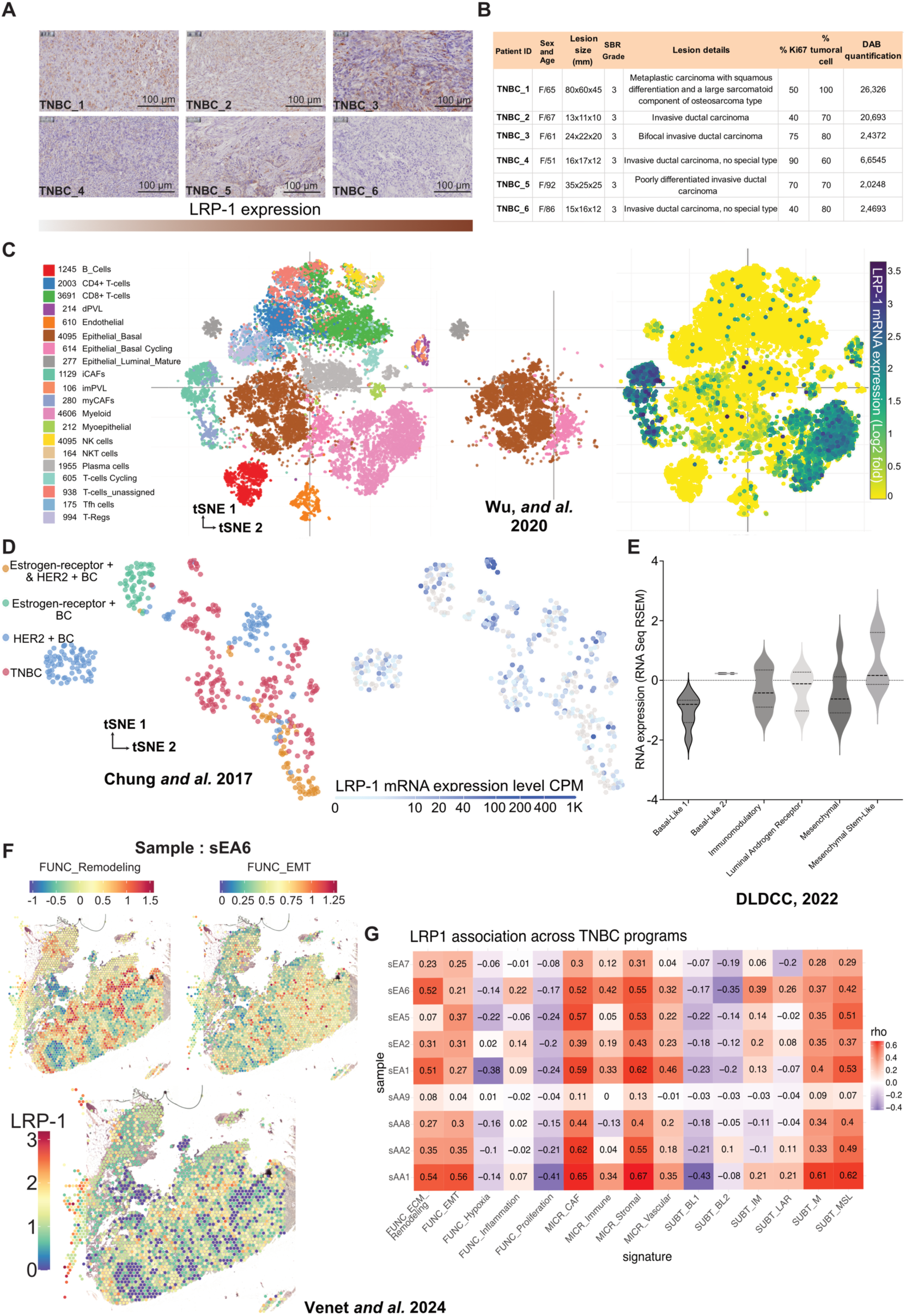
LRP-1 is heterogeneously expressed in TNBC and enriched in mesenchymal /basal epithelial tumor cell populations. **A:** Immunohistochemistry of six high-grade (SBR3) TNBC tumors (Godinot Institute cohort) reveals robust but heterogeneous LRP-1 protein expression, spanning a gradient of DAB staining intensity across diverse histology, including invasive ductal and metaplastic carcinomas. **B:** Clinicopathological annotation confirms cohort heterogeneity despite uniformly high grade. Quantitative DAB analysis shows substantial inter-tumoral variability in LRP-1 levels, independent of parameters such as Ki67 index, lesion size, and tumor cellularity. **C-D:** Single-cell RNA-seq re-analyses demonstrate that LRP-1 is enriched in TNBC relative to other subtypes. In the Wu *and al.* (2020) dataset (n = 24,271 cells), LRP-1 expression localizes preferentially to epithelial basal, myoepithelial, and mesenchymal populations, with minimal expression in immune and luminal compartments. Consistently, re-analysis of the Chung *and al.* (2017) dataset shows higher LRP-1 expression within the TNBC cluster compared to ER+ and HER2+ subtypes. **E:** Bulk transcriptomic analysis from the DLDCC (2022) cohort indicates that LRP-1 is selectively upregulated in the Mesenchymal Stem Like (MSL) TNBC subtype, with lower expression in basal-like (BL1/BL2), luminal androgen receptor (LAR), and immunomodulatory subtypes. **F:** The spatial transcriptomic analysis conducted by Venet *and al*. (2024) on a total of nine triple-negative breast cancer (TNBC) samples reveals that LRP-1 expression correlates with areas of active remodeling and EMT. **G:** Spatial correlation analyses (Venet *and al.*, 2024) link LRP-1 expression to mesenchymal and extracellular matrix (ECM) remodeling programs, with inverse associations to immune and proliferative signatures.

Given the cellular heterogeneity observed histologically, we next interrogated LRP-1 expression at single-cell resolution. Re-analysis of the Chung *and al.*[29] single-cell transcriptomic dataset confirmed that *LRP-1* mRNA is markedly enriched within the TNBC cellular cluster relative to ER+ and HER2+ breast cancer subtypes (Fig. 1D). Further deconvolution of the TNBC tumor microenvironment using the Wu *and al.*[30] scRNA-seq dataset (n = 24,271 cells) mapped preferential LRP-1 expression to epithelial basal, myoepithelial, and mesenchymal cell populations (Fig. 1C). In agreement with these single-cell topological findings, bulk transcriptomic analysis of the DLDCC (2022)[31] patient cohort demonstrated that LRP-1 expression is significantly elevated specifically within the Mesenchymal Stem Like (MSL) TNBC molecular subtype, diverging from basal-like (BL1/BL2), luminal androgen receptor (LAR), and immunomodulatory subtypes (Fig. 1E).

To further characterize the spatial organization and functional networks associated with LRP-1 upregulation, we mapped its expression using spatial transcriptomics (ST) across nine TNBC tissue sections[32]. Superimposition of LRP-1 spatial gene expression onto matched Hematoxylin Eosin (H&E) histology revealed distinct regional concentration, with transcripts heavily enriched in infiltrating tumor zones and stromal-adjacent interfaces (Fig. 1F, fig_additional_1 and fig_additional_2). Crucially, spatial correlation analysis against established TNBC transcriptional programs revealed a robust positive association between localized LRP-1 expression and spatial signatures driving ECM remodeling and Epithelial-Mesenchymal Transition (EMT) (Fig. 1G). Conversely, LRP-1 expression demonstrated inverse correlations with spatially defined immune infiltration and proliferative signatures. Together, these multi-cohort and multi-omic analyses reveal marked heterogeneity of LRP-1 expression across TNBC. LRP-1 is enriched in basal, myoepithelial, and mesenchymal cell populations and is preferentially expressed in the MSL molecular subtype. Spatial transcriptomic analyses further associate LRP-1-rich regions with ECM-remodeling and EMT-related transcriptional programs. Although these observations remain correlative, they provide a rationale for investigating the functional contribution of tumor-cell LRP-1 to TNBC progression and tumor microenvironment remodeling.

### LRP-1 orchestrates proteomic and secretomic remodeling in TNBC Cells

To explore the tumoral role of LRP-1, we generated stable Knock-out (KO)-LRP-1 models of human (HS578-T) and murine (4T1) TNBC cells using CRISPR/Cas9 (KO-LRP-1) (fig_additional_3). In order to define the way and molecular mediators of LRP-1’s tumoral activity, we performed parallel label-free secretomic (Fig. 2A, fig_additional_4) and proteomic profiling (fig_additional_4, fig_additional_5A) between 4T1 WT and KO-LRP-1 cells lines. At the proteome level, the dataset indicates modulation of 133 proteins (reported as 73 up-regulated and 60 down-regulated) (fig_additional_5B), while the secretome exhibited larger remodeling with 345 proteins significantly altered (178 up-regulated and 167 down-regulated) (fig 2B). Pathway and Gene Set Enrichment Analyses performed across multiple annotation resources (Gene Ontology, Hallmark, Reactome and Ingenuity Pathway Analysis), and thanks to using SRplot, showed that the differentially regulated proteins cluster into biologically coherent groups (fig 2C, 2D, fig_additional_5C, 5D, 5E, fig_Additional_6A, 6B). A leading theme common to both datasets is the regulation of proteins implicated in epithelial cell migration and invasion. Representative proteome hits include ETS1, P2RX4, CCBE1, THBS1 and NR2F2 (fig_additional_5F), while the secretome contains several secreted effectors linked to motility and guidance such as FGFBP1, NRP1, SEMA5A, ENPP2, and SLIT2 (Fig. 2A, 2D). The proteomic changes show altered intracellular trafficking, secretion and ECM organization (fig_additional_5F). Notable examples include components and regulators of TGF-β response (e.g. THBS1, LTBP3, SMAD5) (fig_additional_5F), clathrin-coated vesicle pathways (e.g. AP1M2, RGS19) (fig_additional_5F), secretory granules (e.g. RAB3D) (fig_additional_5F), and proteins involved in ECM organization and organelle membrane composition (e.g. CCN2, APLP2, IMMT) (fig_additional_5F). LRP-1 loss induced particularly extensive remodeling of the secreted proteome. Most important affected themes were epithelia mesenchymal transition (EMT) signatures and multiple layers of ECM biology (fig 2D). EMT-related changes included both matricellular factors and signaling molecules (for example, CCN1/2, FN1, POSTN, SMAD3, WNT10A) (fig 2D). ECM remodeling included enzymes and pathways involved in glycosaminoglycan metabolism, matrix degradation and collagen biosynthesis/processing, collectively pointing to large-scale changes in matrix composition and maturation (fig 2D, fig_Additional_7). Consistent with matrix remodeling, proteins associated with the plasma membrane and cell–matrix interactions were strongly affected. Enrichment analyses highlight integrin signaling and integrin ECM interactions, adherens junction pathways and an assortment of membrane receptors and transporters altered upon LRP-1 ablation (representative examples include FN1, COL4A1/4A5, NRP1, FLT1 and multiple SEMA family members) (fig 2D, fig_Additional_8). Finally, the secretome changes reveal substantial modulation of immune and inflammatory networks. LRP-1 depletion was associated with signatures of interferon responses (IFN-γ/α), TNF-α/NF-κB signaling, a broad array of interleukin cascades (including IL-1, IL-6, IL-10, IL-12, IL-13, IL-15, IL-17 and others), complement and acute-phase responses, neutrophil degranulation and antigen-processing pathways. Representative immune-related proteins altered in the secretome include CXCL10, PSMB9, ISG15, CCL5, MYD88 and TYK2 (fig 2D, fig_Additional_9).

**Figure 2:**
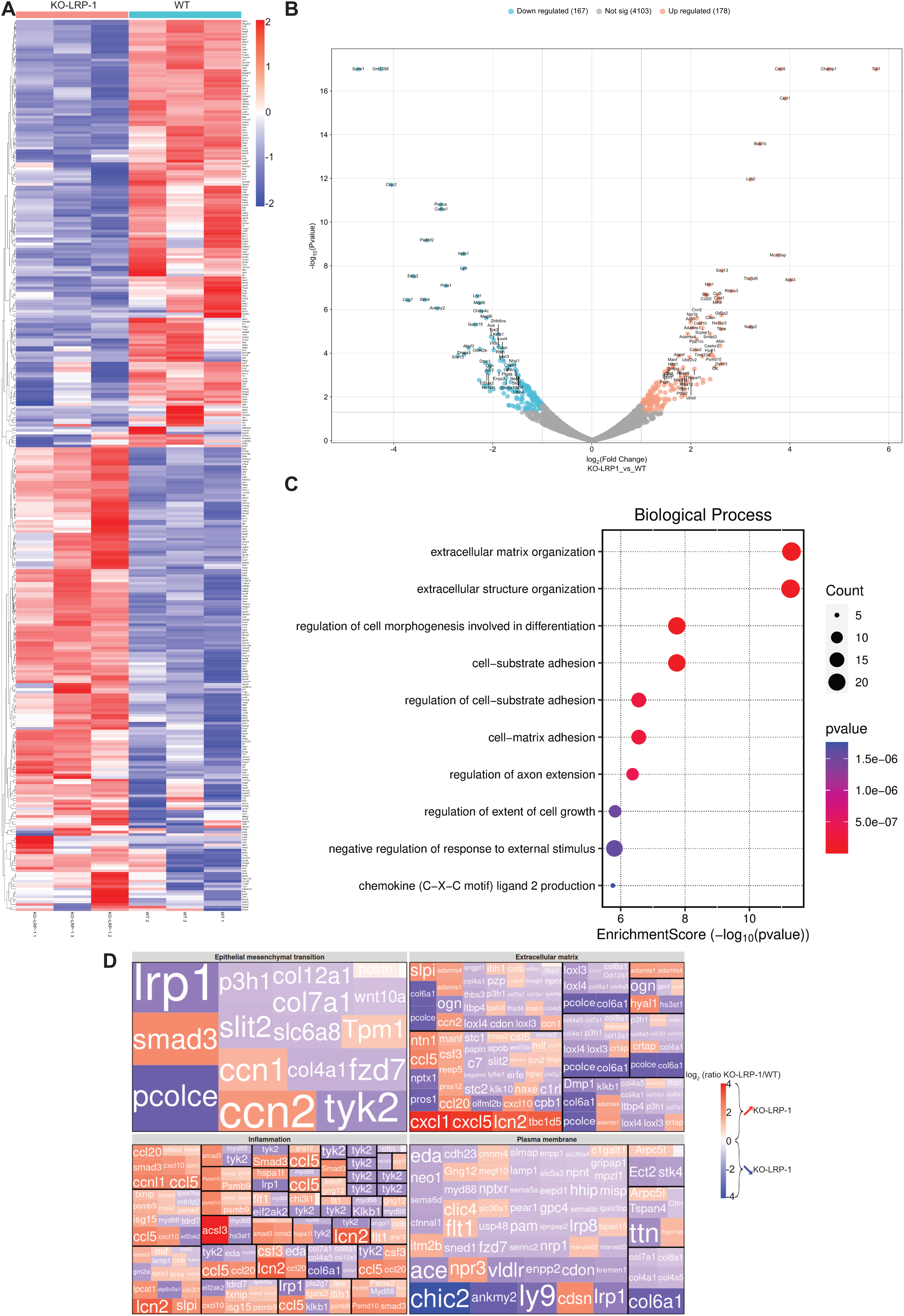
LRP-1 drives secretomic rewiring and pathway-level changes. Overview of panels showing **A:** Heatmap of 345 proteins differentially expressed between 4T1 WT and 4T1 KO-LRP-1 cells. Values are logarithmic scale of fold change from −2 to 2, blue = decreased, red = increased. **B:** Volcano plot of differential protein expression between WT and LRP-1 KO cells X = log2 fold change, Y= −log10 p-value; 167 proteins significantly down-regulated and 178 significantly up-regulated in KO (significant points colored). **C:** Dot plot of S-scores for enriched biological processes. Point color indicates the p-value and point size indicates the number of LRP-1–regulated proteins that contribute to each term. **D:** Treemap of GSEA-derived biological processes impacted by LRP-1 loss (analyses performed across Hallmark, GOCC and IPA). The map is organized into four major process clusters : epithelial–mesenchymal transition (EMT), extracellular matrix (ECM), plasma membrane and inflammation, each containing constituent pathways. Color shows the median log₂ (KO/WT) ratio for those proteins (scale −4 → +4; blue = lower in KO, red = higher in KO). Some pathways are grouped into sub-clusters outlined with black borders; full term-level details are provided in Supplementary_Figure. Only terms meeting significance thresholds are displayed. proteomic figures shown here come from for three independent experiments following total peptide quantity normalization. Protein identifications were retained at an FDR of <1%, and missing values were imputed using the Low Abundance Resampling approach. Differential abundance was defined by P < 0.05 and a fold change of ≤0.5 or ≥2.0.

Together, these data indicate that loss of LRP-1 in 4T1 cells coordinated shifts in proteostasis, ECM composition and immune related signaling. Such multi layered changes are likely to reshape tumor cell behaviour and the tumor microenvironment, with potential consequences for invasion, immune interactions and matrix remodeling. Even if these targets are identified in the mouse model, functional tests and complementary molecular analyses will be conducted across species.

### LRP-1 expression in TNBC tumors promotes both 2D and 3D migration and invasion

To explore the functional role of LRP-1, we performed numerous migration and invasion assays. In wound healing assays (Fig. 3A), loss of LRP-1 significantly reduced migratory capacity in both TNBC cell lines. In HS578-T cells, wound closure dropped from 96.17% in control conditions to 62.06% in knockout cells. A similar reduction was observed in 4T1 cells, where closure decreased from 90,80% to 47,27% (Fig. 3A). Moreover, 24-hour tracking single-cell migration monitoring demonstrated that KO-LRP-1 drastically limits cellular reach and dispersion. Radar tracking and euclidean distance quantification confirmed this impairment, with KO-LRP-1 cells showing reduced motility in both 4T1 and HS578-T lines on fibronectin (∼61–76% decrease respectively) and collagen (∼50% decrease) coating (Fig. 3B). Because two-dimensional assays do not fully reflect the complexity of the tumor progression, we next turned to a three-dimensional collagen gel migration model. Live-cell videomicroscopy over 24 hours revealed striking differences in cell behavior : trajectories of KO-LRP-1 cells were noticeably shorter and less dynamic compared to controls (Fig. 3C). Quantitative measurements confirmed a significant reduction in migration speed in both cell lines. In HS578-T cells, median migration speed fell from 14.5 µm/h with LRP-1 to 4.69 µm/h after knockout. In 4T1 cells, speed declined from 16.71 µm/h in WT cells to 4.19 µm/h in knock-out cells (Fig. 3C). To determine whether reduced migratory capacity was accompanied by impaired invasiveness, we performed modified Boyden chamber assay. KO-LRP-1 cells exhibited significantly reduced invasive potential compared with WT cells. In HS578-T KO-LRP-1 cells, invasion dropped by 36.27%, while in 4T1 KO-LRP-1 cells the effect was more marked, with a reduction of 74.51% (Fig. 3D). Together, these results demonstrate that LRP-1 depletion markedly impairs migratory and invasive capacities of TNBC cells across complementary 2D and 3D experimental models.

**Figure 3:**
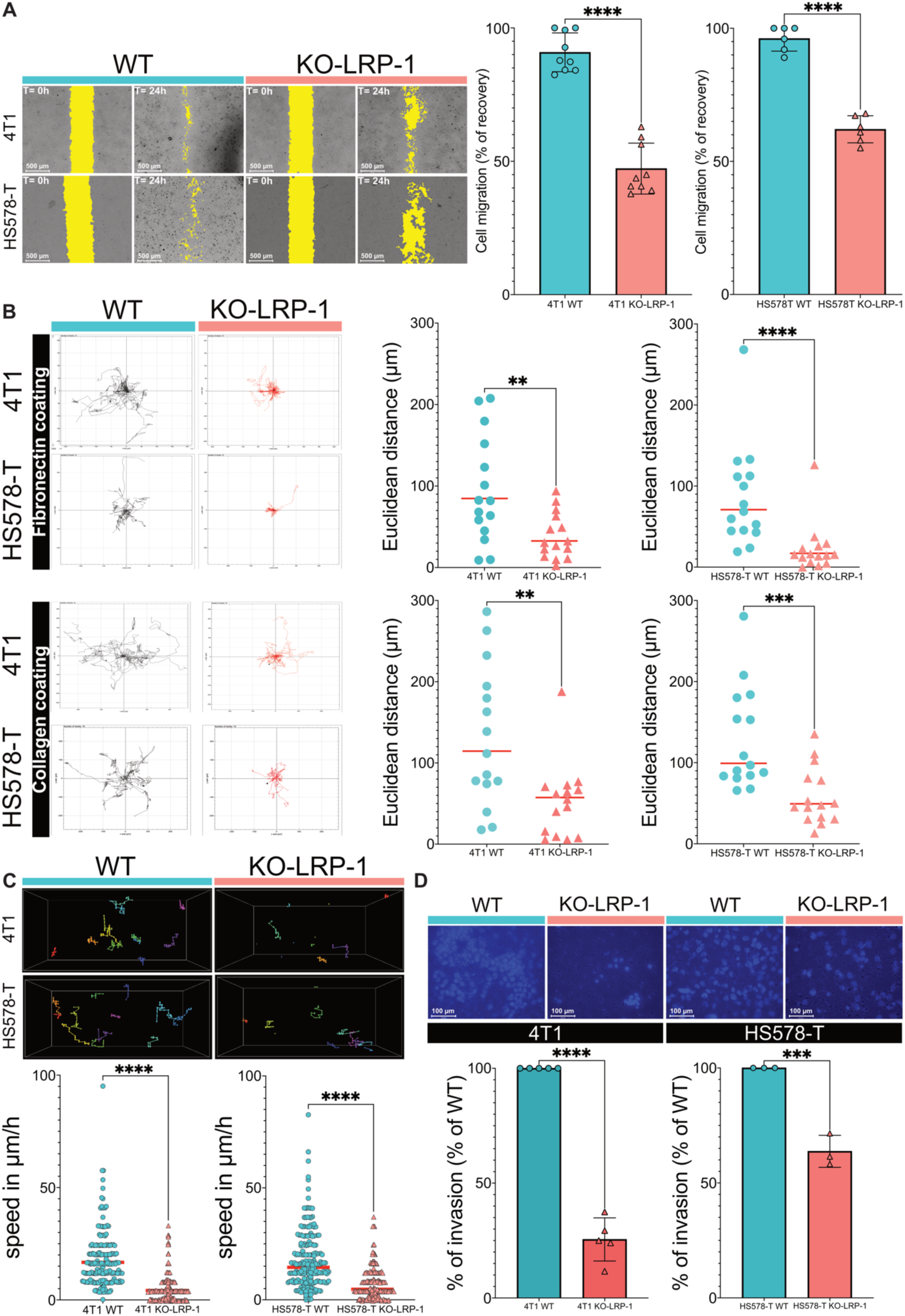
LRP-1 deficiency suppresses collective, individual, and 3D migration and invasive capacity. **A:** Collective cell migration assessed by wound healing assay. Representative images highlight the wound area at T0 and closure progress. Bar graphs quantify the significant impairment of wound closure in LRP-1-knockout (KO-LRP-1) 4T1 and HS578-T cells compared to wild-type (WT) controls. Data are presented as mean ± SD based on data from three independent experiments. Statistical analysis was performed using an unpaired two-tailed Student’s *t*-test. **B:** Monodispersed migration on fibronectin or collagen-coated surfaces. Representative radar plots illustrate individual cell trajectories from a common origin. Quantitative analysis of displacement parameters (e.g. Euclidean distance) reveals a marked reduction in single-cell motility upon LRP-1 loss across different extracellular matrix (ECM) substrates. Data are presented as scatter dot plots with the median based on data from 15 independent cells for each independent cell line. Statistical analysis was performed using the two-tailed Mann–Whitney test. **C:** 3D migration in collagen gels. Representative 3D tracks and quantification of migration speed indicate that LRP-1 depletion restricts the ability of cancer cells to navigate through a complex 3D matrix. Data are presented as scatter dot plots with the median based on data from three independent experiments. Statistical analysis was performed using the two-tailed Mann–Whitney test. **D:** Modified boyden chamber assay. Representative images of DAPI stained nuclei show cells that successfully invaded through matrigel coated membranes. Quantitative analysis confirms a large loss of invasive capacity in KO-LRP-1 cells. Data are shown as bar graphs and presented as mean ± SD based on data from three (HS578-T WT/KO-LRP-1) or five (4T1 WT/KO-LRP-1) independent experiments. Statistical analysis was performed using an unpaired two-tailed Student’s *t*-test. For all graphs, statistical significance is indicated as follows: *P < 0.01 (*\*\*), *P* < 0.001 (***), and *P* < 0.0001 (****).

### LRP-1 controls TNBC spheroid morphogenesis and invasive architecture

To further assess the impact of LRP-1 on tumor growth and invasion, we perform tumor spheroid model. Loss of LRP-1 markedly impaired both spheroid growth and invasion in triple negative breast cancer cells tumor spheroids. As shown in Fig. 4A, 4T1 and HS578-T KO-LRP-1 cells formed significantly bigger spheroids compared to their wild-type counterparts at day 14 (0,76 mm^2^and 0,96 mm^2^ in 4T1 WT and 4T1 KO respectively, 0,75 mm^2^ and 0,81 mm^2^ in HS578-T WT and HS578-T KO respectively) (Fig. 4A). Moreover, in a 3D matrix, WT spheroids displayed numerous evasive structures and an irregular invasive front, whereas KO-LRP-1 spheroids showed a compact morphology with markedly reduced invasive area (−78% for 4T1, - 49% for HS578-T). These data indicate that LRP-1 promotes spheroid invasion in this triple negative breast cancer model (Fig. 4B).

**Figure 4:**
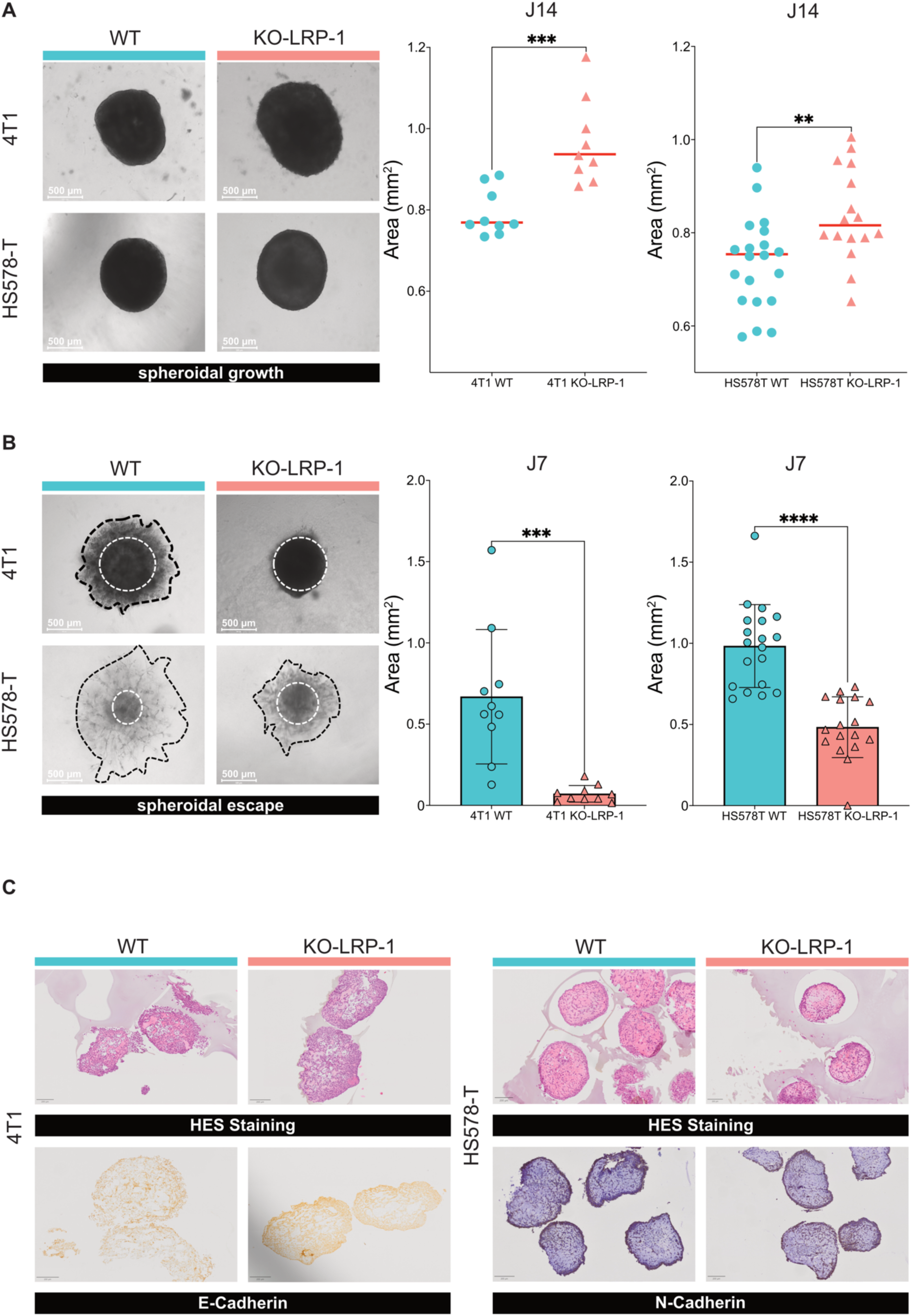
LRP-1 deficiency promotes spheroid growth while inhibiting invasive escape and modulating cadherin expression. **A:** 3D growth of 4T1 and HS578-T spheroids. Representative bright-field images and quantification of the spheroid area at day 14 (D14) show a significant increase in size for LRP-1-knockout (KO-LRP-1) spheroids compared to wild-type (WT) control spheroids. Data are presented as scatter dot plots with the median indicated based on data from three independent experiments. Statistical analysis was performed using an unpaired two-tailed Student’s *t*-test. **B:**Spheroid escape assay evaluating invasive potential in a 3D matrix. Representative images illustrate the spheroid core (white dashed circle) and the invasive outgrowth (black dashed line). Quantification at day 7 (D7) demonstrates that LRP-1 depletion significantly impairs the invasive escape area in both mammary spheroids. Data are shown as bar graphs and presented as mean ± SD based on data from three independent experiments. Statistical analysis was performed using an unpaired two-tailed Student’s *t*-test. **C:** Histological and phenotypic characterization of spheroids. Hematoxylin-Eosin-Saffron (HES) staining shows the internal structural morphology. Immunohistochemical (IHC) analysis highlights the expression levels and distribution of E-cadherin (in 4T1) and N-cadherin (in HS578-T) in WT versus KO-LRP-1 spheroids, reflecting alterations in cell-cell adhesion markers. Data are presented as representative images performed on 12 independent spheroids. For all graphs, statistical significance is indicated as follows: *P < 0.01 (*\*\*), *P* < 0.001 (***), and *P* < 0.0001 (****).

A structural characterization of the spheroids was initiated to better assess their architectural organization. Spheroids were stained with H&E to evaluate overall morphology and the extent of necrotic areas. Morphological analysis revealed that 4T1 and HS578-T WT spheroids displayed a peripheral crown of cohesive cells surrounding a central necrotic core. KO-LRP-1 spheroids exhibit a markedly altered morphology, showing a fragmented island-like organization. These morphological differences suggest that LRP-1 loss impacts spheroid organization. Upon LRP-1 depletion, E-cadherin staining appeared increased in 4T1 spheroids, whereas N-cadherin staining was reduced in HS578-T spheroids, consistent with attenuation of EMT-associated features (Fig. 4C). Together, these results show that LRP-1 contributes to the invasive architecture of TNBC spheroids and influences their structural and cadherin-associated organization.

### LRP-1 shapes the molecular and mechanical basis of TNBC cell invasiveness

To further characterize the effect of LRP-1 on the migratory capacity of TNBC cancer cells, anti-vinculin immunofluorescence and microfilament labeling experiments were performed. For 4T1 cell lines, WT cells adopt a migratory phenotype represented by a stretched morphology, a developed actin cytoskeleton and numerous focal adhesion contacts at the migratory front and retracted queue. HS578-T WT cells exhibit large polarized lamellipodia and focal adhesion complexes at the leading edge in accordance with a migratory phenotype. In both cell line, knockout of LRP-1 induces profound cell shape remodeling In 4T1. KO-LRP-1 affects cell spreading and a decrease of cell focal adhesion complexes. In HS578-T, KO-LRP-1 is associated with a round phenotype and numerous focal adhesion complexes all around cells. These morphological changes are consistent with our previous observations concerning the effect of LRP-1 on cell migration (Fig. 5A).

**Figure 5:**
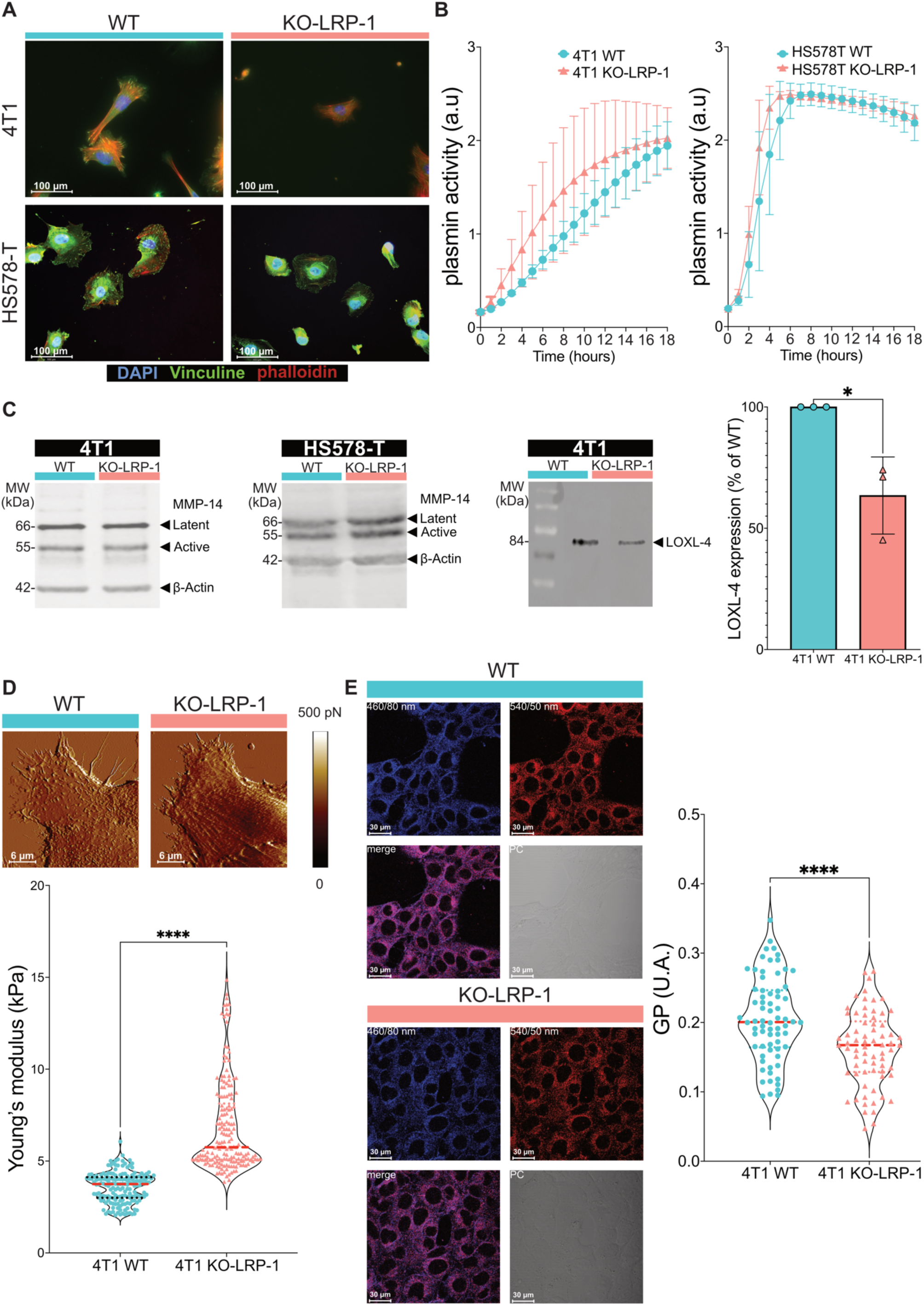
LRP-1 KO remodels cytoskeletal organization, extracellular matrix, and cell mechanics. **A:** Representative immunofluorescence images of WT and KO-LRP-1 4T1 and HS578T cells stained for F-actin (phalloidin, red), vinculin (green), and nuclei (DAPI, blue). Images illustrate changes in cell morphology, actin cytoskeleton organization, and focal adhesion distribution upon LRP-1 loss. **B:** Time-course analysis of plasminogen activation in 4T1 (left) and HS578T (right) WT and KO-LRP-1 cells over 18 h. Data are presented as mean ± SD based on data from three independent experiments. **C:** Representative immunoblots showing latent and active MMP-14 expression in 4T1 and HS578T WT and KO-LRP-1 cells, and LOXL-4 expression in 4T1 cells. Relative LOXL-4 protein levels are shown as bar graphs and presented as mean ± SD from three independent experiments. Statistical analysis was performed using an unpaired two-tailed Student’s *t*-test. **D:** Cell stiffness was assessed by atomic force microscopy (AFM) in 4T1 WT and KO-LRP-1 cells. The violin plot represents the distribution of individual young’s modulus measurements with median and quartiles indicated based on data from four independent experiments. Representative image are shown on the top. Statistical analysis was performed using the two-tailed Mann–Whitney test. **E:** Membrane lipid order was evaluated in 4T1 WT and KO-LRP-1 cells by Laurdan (calculated generalized polarization (GP)) imaging using the indicated emission channel. Representative fluorescence, merged, and phase-contrast images are shown. The violin plot represents the distribution of individual GP values, with median and quartiles indicated based on data from seventy independent cells to each cell line. Statistical analysis was performed using the two-tailed Mann–Whitney test. For all graphs, statistical significance is indicated as follows: *P* < 0.0001 (****).

To determine whether cellular invasion in TNBC cell lines is mediated in part by proteolysis, we studied plasminogen activator (PA) activity and invasion-associated enzymes expression. Loss of LRP-1 doesn’t significantly impaired PA activities neither in 4T1 nor in HS578-T cells (Fig. 5B). MMP-14 expression is not altered neither in both cell lines by KO-LRP-1 (Fig. 5C). However, the 4T1 aggressive cell line have shown a reduction of LOXL-4 enzyme expression in LRP-1 knockouts compared to the WT cells by approximately 36.5% (Fig. 5C). These findings suggest a potential association between LRP-1 expression and LOXL-4 abundance. To examine whether the LOXL-4 reduction was accompanied by changes in the cells’ mechanical and physico-chemical properties, we measured cell stiffness by atomic force microscopy (AFM) and assessed membrane order using Laurdan imaging. AFM peak-force measurements showed that the mean Young’s modulus increased from 3.57 kPa in LRP-1–expressing cells to 6.84 kPa in KO-LRP-1 (Δ = 3.27 kPa) (Fig. 5D). Laurdan-based GP measurements indicated a decrease in lipid ordering in KO-LRP-1 cells (shift toward Ld domains) (Fig. 5E). Altogether these data show that loss of LRP-1 is not accompanied by reduction of MT1-MMP but coincides with reduced LOXL-4 and increased membrane stiffness. This pattern suggests that the impaired invasiveness observed upon LRP-1 depletion stems primarily from altered matrix remodeling and changes in the cells’ mechanical/physico-chemical properties rather than from direct upregulation of metalloproteinases.

### LRP-1 drives TNBC tumor growth and immune microenvironment remodeling *in vivo*

In order to evaluate the effect of LRP-1 on tumor growth, 4T1 cells (5×10^5^) were injected in mammary fat pad of syngenic Balb/C mice. Tumor growth was followed for 26 days and results demonstrate a significant decrease of tumor volume in mice bearing 4T1 KO-LRP-1 tumors (mean size 357 mm^3^ *vs* 111 mm^3^) (Fig. 6A/6B). After excision, tumors were stained by Masson’s trichrome and collagen quantification was obtained by using Image J color deconvolution2 function. Red channel intensity quantification after thresholder revealed a 37.6% decrease in collagen deposition in 4T1 KO-LRP-1 tumors (Fig. 6C). Variation of collagen deposition seems to be localized at the tumor periphery. This observation was further confirmed by SHG imaging, which demonstrated a clear decrease in SHG-positive fibrillar collagen signal in 4T1 KO-LRP-1 tumors compared with 4T1 WT tumors (Fig. 6D).

**Figure 6:**
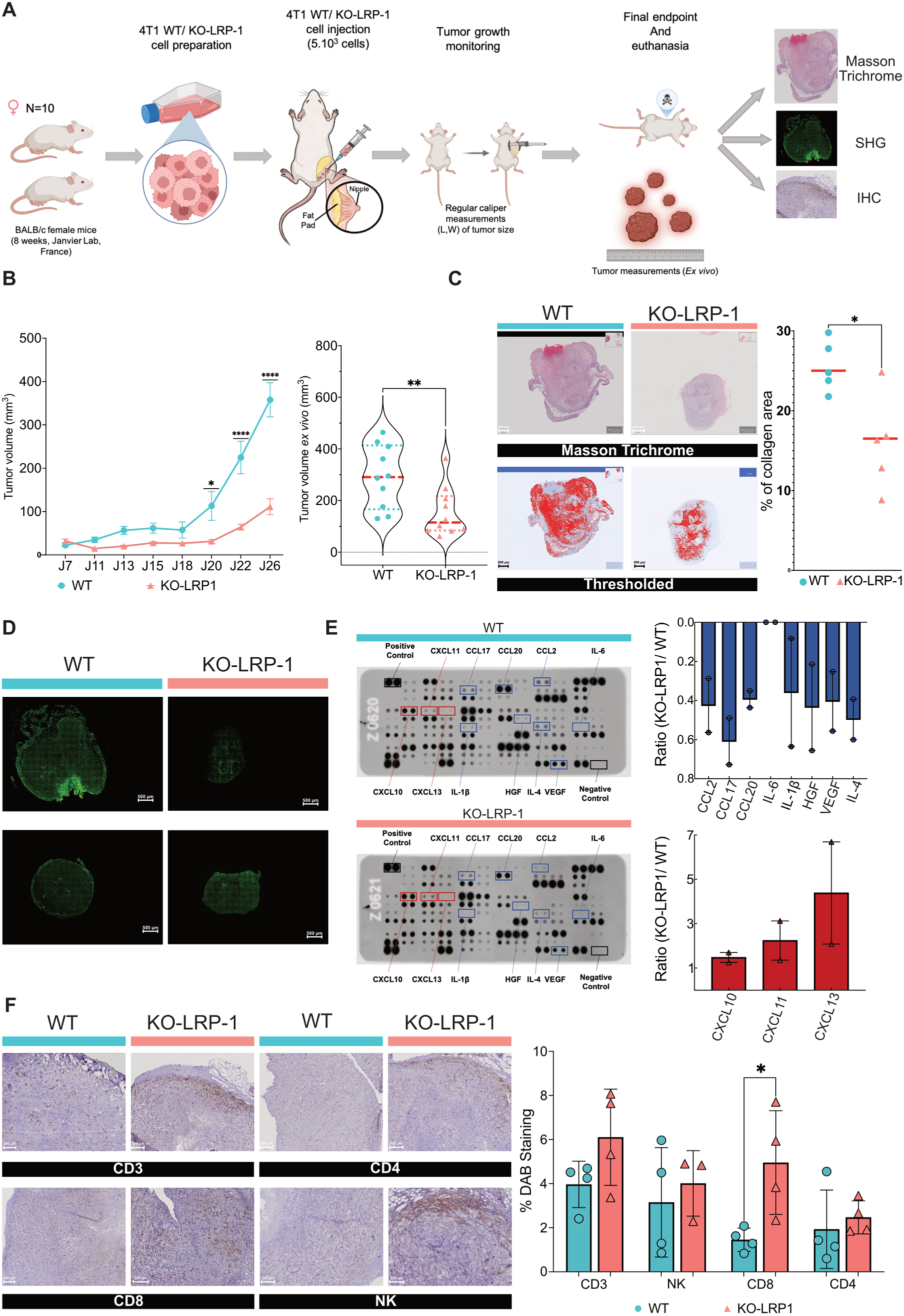
LRP-1 deficiency suppresses mammary tumor growth and modulates the inflammatory microenvironment. **A**: Experimental workflow : 5.10^5^ cells (4T1 WT or KO-LRP-1) were orthotopically injected into BALB/C mice, followed by longitudinal monitoring and terminal histological analysis. **B:** Tumor growth kinetics (left) and terminal *ex vivo* tumor volume measurements (right), showing significantly reduced progression in KO-LRP-1 mice. Data are presented as mean ± SEM (n= 10 mice per group) and the violin plots represent the distribution of individual terminal *ex vivo* tumor volumes, with median and quartiles. Statistical analysis was performed using a two-way ANOVA test for growth kinetics and two-tailed Mann–Whitney test for terminal *ex vivo* tumor volume comparisons. **C:** Representative Masson’s Trichrome stainings (top, collagen : blue channels) and thresholded quantification (bottom, collagen : red channels) assessing collagen deposition and fibrosis within the tumor stroma. Data are presented as scatter dot plots with median based on data from three independent experiments. Statistical analysis was performed using two-tailed Mann–Whitney test. **D:** Second Harmonic Generation (SHG) microscopy imaging, confirming a marked reduction in fibrillar Collagen I density in KO-LRP-1 tumors compared to WT. Data are presented as representative image of three independent experiments. **E:** Cytokine array profiling of 4T1 WT and KO-LRP-1 secretome and densitometric quantification of differentially expressed factors. Bar graphs show the ratio of Cytokine/chemokine expression in KO-LRP-1 versus WT tumors and presented as mean ± SEM based on data from two independent experiments. **F:** Immunohistochemical (IHC) staining and quantification of tumor-infiltrating immune cells markers. Data are shown as bar graphs and presented as mean ± SD based on data from three independent experiments. Statistical analysis was performed using an unpaired two-tailed Student’s *t*-test. For all graphs, statistical significance is indicated as follows: *P* < 0.05 (\**), P < 0.01 (*\*\*) and *P* < 0.0001 (****).

We next thought to quantify immune cell accumulation within the tumor microenvironment, hypothesizing that decreased collagen deposition at the tumor periphery in 4T1 KO-LRP-1 tumors could facilitate immune cell infiltration. Consistently, we found an increased accumulation of CD8-positive T cells in 4T1 KO-LRP-1 tumors (Fig. 6F).

Cytokine profiling of the secretomes revealed increased secretion of the chemokines CXCL10, CXCL11, and CXCL13 in the absence of LRP-1, suggesting that LRP-1 invalidation generate a chemokine behavior more permissive to lymphocyte infiltration (Fig. 6E). Conversely, decreased levels of CCL2, CCL17, CCL20, IL-1β, IL-4, IL-6, HGF, and VEGF were detected in KO-LRP-1 secretomes compared with WT cells (Fig. 6E). Together, these results indicate that LRP-1 depletion is associated with altered chemokine and cytokine secretion profiles, reduced collagen deposition, and increased CD8^+^ T-cell accumulation *in vivo*.

## Discussion

Our integrative analyses reveal LRP-1 as a main regulator of TNBC aggressiveness associated with coordinated regulation of tumor cell programs, ECM remodeling, and immune exclusion [16,33,34]. LRP-1 protein is highly expressed yet heterogeneous across TNBC tumors (Fig. 1A–B), localizing to invasive, stromal-adjacent niches. Multi-omic profiling shows that LRP-1 loss in TNBC cells broadly rewires intracellular and secreted proteomes toward reduced EMT/motility, altered ECM and integrin networks, and heightened inflammatory signaling (Fig. 2). Functionally, LRP-1 knockout (KO) impairs cell migration and invasion in 2D and 3D assays (Fig. 3) and yields compact tumor spheroids with blunted invasive fronts (Fig. 4). Mechanistically, LRP-1-deficient cells exhibit dramatic cytoskeletal and adhesion remodeling (Fig. 5A) without upregulating canonical proteases (MMP-2, MMP-9, MMP-14, PA); instead, they show decreased LOXL-4 and increased cortical stiffness (Fig. 5B–D). *In vivo*, KO-LRP-1 TNBC tumors grow poorly, deposit ∼50% less collagen at their invasive margins, recruit more CD8⁺ T cells and NK cells (Fig. 6). The KO-LRP-1 secretome upregulates lymphocyte-associated chemokines (CXCL10/11/13) and downregulates cytokines commonly associated with immunosuppressive or pro-tumor signals (CCL2/17/20, IL-1β/6, VEGF), consistent with a more immune-permissive microenvironment. Together, our data position LRP-1 as a nexus linking integrin signaling, matrix mechanics, and immune infiltration, and suggest that targeting LRP-1 or its downstream effectors could impede TNBC invasion and promote a more immune-responsive tumor microenvironment. Together, our findings identify LRP-1 as a central regulator of ECM remodeling, tumor cell mechanics, and immune microenvironment organization in TNBC.

LRP-1 is a ubiquitously expressed endocytic receptor that integrates matrix and signaling cues [35,36]. Our IHC and single-cell data indicate that high-grade TNBCs can express abundant but heterogeneous LRP-1, especially in regions enriched in mesenchymal-like tumor cells. We confirm prior single-cell profiling [29,30] that LRP-1 mRNA is enriched in TNBC cellular compartments, basal/mesenchymal cells, far more than in luminal or HER2 tumors. Bulk expression data from TNBC cohorts likewise show LRP-1 specifically high in the “Mesenchymal Stem Like” molecular subtype (Fig. 1E), consistent with an association between LRP-1 expression and invasive stromal programs. Recent spatial-transcriptomic mapping [32] detected LRP-1 transcripts in tumor–stroma interfaces and infiltrative fronts (Fig. 1F, fig_additional_1 and fig_additional_2). These high-LRP-1 regions positively track signatures of ECM remodeling and EMT and inversely track immune infiltration programs (Fig. 1G). This aligns with broader TNBC archetype analyses showing that mesenchymal TNBC tumors harbor angiogenic, stromal-rich environments, whereas immune-high subtypes have better prognosis. Thus, LRP-1 is embedded in pro-invasive niches and may contribute to tumor–stroma crosstalk.

LRP-1’s effect on cell migration is multifaceted. Secretomic profiling of KO-LRP-1 cells identifies dozens of altered proteins (Fig. 2A–B), with enrichment for regulators of cell adhesion, trafficking, and TGF-β/ECM pathways (Fig. 2C–D). For example, several known mediators of migration (ETS1, THBS1, NR2F2, CCN2) are dysregulated in KO-LRP-1 cells, and we observe pronounced changes in secreted factors (FGFBP1, NRP1, SEMA5A, SLIT2, etc.) that guide motility. Crucially, integrin-ECM pathways are prominent among affected networks: fibronectins, collagens, plexins/semaphorins, and integrin receptors themselves are altered (Fig. 2D). This resonates with emerging models in which LRP-1 physically binds β1-integrin and its adaptor kindlin-2 to coordinate integrin activation, trafficking, and turnover. Wujak *and al* [37], showed LRP-1 promotes β1 integrin activation (via kindlin-2) and directs integrin to lysosomal degradation [38], ensuring dynamic adhesion assembly and disassembly. In our system, loss of LRP-1 may disrupts this integrin quality-control, potentially altering the surface β1 integrin turnover and directional migration. Indeed, KO-LRP-1 cells show aberrant focal adhesion and actin patterns (Fig. 5A), consistent with defective adhesion cycling seen when LRP-1 is silenced [39].

The loss of LRP-1 profoundly remodels the secretome (345 proteins changed, Fig. 2B) with emphases on ECM and immune pathways (Fig. 2D). Key EMT mediators (matricellular CCN1/2, FN1, POSTN, SMAD3, WNT10A) are dysregulated, and enzymes for glycosaminoglycan synthesis, collagen maturation, and ECM degradation show large shifts. These data suggest that LRP-1 contributes to the regulation of matrix composition and remodeling [25]. This modulation occurs in part through the decrease of ADAMTS-1 expression (Fig. 2A,2D), one known consequence of which is an increased concentration of VEGF in the extracellular environment, capable of inducing tubulogenesis and thereby promoting migratory and invasive processes in TNBC [39]. The results of this study, along with previous findings [16], confirm these data by demonstrating that the loss of LRP-1 leads to both an increase in ADAMTS-1 and a significant disruption of angiogenesis [16]. Other example, KO-LRP-1 cells downregulate LOXL-4 (∼36%), an enzyme that crosslinks collagen and laminin A in the ECM, but also crosslinks annexin A2 on tumor cells and that has also been shown to be an inducer of chemoresistance in TNBC [40]. Moreover, Takahashi *and al*. [41] recently showed that TNBC cells secrete LOXL-4 to multimerize annexin A2 on their surface, which binds β1 integrin and prevents integrin internalization. This creates a feed-forward circuit: LOXL-4 keeps integrins at the membrane to sustain invasion. In our data, LRP-1 loss reduces LOXL-4, which may weaken this pro-invasive circuit and potentially affect integrin-dependent adhesion dynamics. Thus, LRP-1 may indirectly contribute to integrin-dependent invasion through maintenance of the LOXL-4/annexin A2 signaling axis. Notably, despite this, classic proteolytic pathways do not compensate: plasmin activity levels are unchanged with KO-LRP-1, suggesting invasion defects are independent of canonical proteases (MMP-2, MMP-9, MMP-14, PA), as we also evidenced in a past study [42].

Besides, LRP-1 knock-out substantially remodels immune-related secretions. We find increased IFN signatures and cytokines (CXCL10/11, IL-12, complement factors) alongside reduced immunosuppressive factors (CCL2, IL-6, VEGF) (Fig. 2D). These changes are consistent with a tumor-associated secretory profile that may favor lymphocyte recruitment, in agreement with the increased CD8^+^ T-cell accumulation and the trend toward higher NK-cell infiltration observed in KO-LRP-1 tumors.

The functional impact of these molecular changes is consistent: with major impairment of cell motility upon LRP-1 knockout. In 2D wound-healing and single-cell tracking, KO-LRP-1 TNBC cells are dramatically less migratory (Fig. 3A–B). This replicates earlier findings in breast cancer models [42] that LRP-1 loss stalls invasion even when proteases rise. In 3D collagen assays, KO-LRP-1 cell trajectories are shorter and slower (Fig. 3C), pointing to impaired deformability or traction. Correspondingly, transwell invasion through matrigel drops ∼36–75% upon LRP-1 loss (Fig. 3D). These concordant 2D and 3D defects underscore LRP-1’s role in sustaining the migratory/invasive phenotype of TNBC.

At the tissue scale, TNBC spheroids lacking LRP-1 also show altered behavior (Fig. 4). Interestingly, KO-LRP-1 spheroids grow slightly larger in area than controls by day 14 (Fig. 4A), yet their invasive projections are almost abolished (– 78% area for 4T1, – 49% for HS578-T, Fig. 4B). Histologically, WT spheroids form a peripheral cohesive shell around necrotic cores, whereas KO spheroids fragment into dense, “island-like” clusters (Fig. 4C). One possible interpretation is that LRP-1 loss may uncouple spheroid expansion from invasive abilities, leading to tighter spheroids that expand but fail to sprout. The mechanisms underlying the increased spheroid area remain unclear and may involve alterations in spheroid architecture and cell organization rather than changes in invasive behavior alone. Changes in cadherin staining patterns further indicate altered cell–cell adhesion organization and reduced mesenchymal features following LRP-1 depletion. Collectively, these findings indicate that LRP-1 regulates not only single-cell motility but also the invasive architecture of multicellular TNBC spheroids.

LRP-1 loss also remodels the internal machinery of invasion. Immunofluorescence imaging shows that WT TNBC cells display classic migratory polarity: elongated shape, polarized lamellipodia, and trailing focal adhesions (Fig. 5A). In contrast, KO-LRP-1 cells lose this polarity. In 4T1, cells are less spread with fewer front adhesion complexes, in HS578-T they become round with adhesions encircling the cortex (Fig. 5A). These divergent morphologies suggest LRP-1’s integration with integrin and cytoskeletal regulators. Indeed, LRP-1 interacts with kinases (ERK/JNK) and proteases (calpain) [43] to modulate adhesion turnover, so its loss could shift the balance of adhesion assembly/disassembly [44]. The net effect is defective front–rear coordination that stalls migration despite ample adhesion formation.

Mechanical measurements corroborate these structural changes [45]. Atomic force microscopy shows KO-LRP-1 cells are significantly stiffer (mean Young’s modulus ∼6.8 vs 3.6 kPa, Fig. 5D). A stiffer actin cortex may reduce cell deformability through dense matrices, These data support a broader mechano-phenotypic shift under LRP-1 extinction compatible with an altered invasive phenotype. Intriguingly, our Laurdan lipid imaging (Fig. 5E) indicated KO-LRP-1 membranes are more fluid, revealing a “stiffness–fluidity” disconnect: likely, the cortex’s actin dominates AFM stiffness, while lipid packing changes reflect separate lipidomic effects of LRP-1 (e.g. altered cholesterol homeostasis [46]). Regardless, the mechanical shift underscores that LRP-1 tunes the cell’s physical phenotype, consistent with its role as a mechanochemical integrator at adhesion sites.

The *in vivo* data integrate these findings: 4T1 tumors lacking LRP-1 grew much more slowly (Fig. 6A–B), contrasting with the larger projected area observed in spheroids, suggesting that stromal, immune and angiogenic components [16] of the *in vivo* microenvironment may critically influence the tumor phenotype. Upon histological and SHG analyses, KO-LRP-1 tumors exhibited markedly reduced collagen deposition and fibrillar collagen signal (Fig. 6C–D). This is significant because desmoplastic collagen at tumor margins can stiffens the niche and support invasion. In TNBC and other cancers, high collagen density has been associated with impaired T cells infiltration and may also signals via collagen receptors such as LAIR-1 to suppress immunity. Consistent with this, KO-LRP-1 tumors showed markedly more CD8⁺ T cells and CD4⁺ T/NK cells (Fig. 6F), suggesting that tumor-cell LRP-1 may contribute to an immunosuppressive microenvironment. This is in line with prior evidence linking collagen-rich stroma to reduced lymphocyte infiltration and impaired response to immunotherapy.

Cytokine profiling clarifies mechanisms (Fig. 6E): KO-LRP-1 cells secreted higher levels of CXCL10, CXCL11, and CXCL13, chemokines known to attract and organize T/B cells. CXCL10/11 engage CXCR3 on effector T cells and NK cells (often IFN-induced), while CXCL13 recruits B cells and T follicular helper cells to form tertiary lymphoid structures. Indeed, elevated CXCL13 has been linked to TLS formation and better breast cancer prognosis. Meanwhile, KO-LRP-1 dampened secretion of CCL2, CCL17, CCL20, IL-1β, IL-4, IL-6, HGF, and VEGF factors, frequently associated with pro-tumor inflammation, myeloid-cell recruitment or immune suppression [47]. Notably, Mariathasan *and al.* [48] showed that stromal TGF-β drives collagen deposition and blocks T-cell infiltration, and that blocking TGF-β reduces collagen I production and permits T cells to penetrate tumors. Our findings are consistent with this conceptual framework : by disrupting LRP-1, we possibly blunt a collagen/TGF-β axis and remodel the chemokine milieu to favor immune attack.

## Conclusions

In summary, our data support a model in which LRP-1 acts as a molecular hub in TNBC, linking integrin-mediated adhesion, matrix remodeling, and immune exclusion to drive tumor progression. LRP-1 sustains aggressive mesenchymal phenotypes by modulating extracellular matrix remodeling, and inflammation-associated secreted factors and by promoting a pro-invasive secretory program including LOXL-4-mediated matrix crosslinking. Loss of LRP-1 disrupts this network: cells stiffen and lose motility, collagen deposition decrease, and immune infiltrates increase. These insights suggest new therapeutic angles. Inhibiting LRP-1 or its downstream signaling could help reduce matrix-associated barriers and promote a more immune-infiltrated tumor microenvironment.. Interestingly, biomarkers of LRP-1 expression or activity may help identify ECM-rich TNBC subsets to benefit to future stroma-targeting therapies.

## Acknowledgement

The authors thank Dr Christine Terryn of URCAtech PICT platform for technical assistance in confocal microscopy and SHG analyses.

The authors thank Nathalie Lalun and Nicole Bouland of URCAtech Phim platform for technical assistance in immunohistochemistry experiments.

## Author information

Maxence Mocquery-Corre is the first author

Lucille Cartier and Abdel-Ilah Aziz are co-second author and contributed equally to this work. Jérôme Devy and Jessica Thevenard Devy are co-last author and contributed equally to this work.

## Authors and Affiliations

**Matrice Extracellulaire et Dynamique Cellulaire - UMR CNRS 7369 – Université de Reims Champagne-Ardenne, Centre National de la Recherche Scientifique – France**

Maxence Mocquery-Corre, Abdel-Ilah Aziz, Alexandre Berquand, Julie Clachet, Chloé Jean, Hassan El Btaouri, Cathy Hachet, Lise Chazée, Katia Savary, Camille Boulagnon Rombi, Stéphane Dedieu, Jérôme Devy & Jessica Thevenard-Devy

**Département de Recherche, Institut Godinot, UR7509 IRMAIC Reims, France**

Lucille Cartier, Yacine Merrouche & Stéphane Potteaux

**faculty of medical sciences, Faculty of Medical Sciences, UM6P Hospitals, University** Mohammed VI Polytechnic, Benguerir, Morocco

Abdel-Ilah Aziz

**Centre de Recherche en Sciences et Technologies de l’Information et de la Communication, EA 3804, Université de Reims Champagne-Ardenne, France**

Chloé Jean

**Univ. Bordeaux, CNRS, INSERM, TBM-Core, US5, UAR 3427, OncoProt, F-33000 Bordeaux, France**

Anne-Aurélie Raymond & Jean-William Dupuy

**Centre de ressources biologiques, Institut Godinot, Reims, France.**

Célia Maquin & Eva Brabencova

**CNRS UMR 7039 CRAN, Université de Lorraine, Vandœuvre-lès-Nancy, France.**

Muriel Barberi-Heyob

**Université de Reims Champagne Ardenne, IRMAIC UR 7509, Reims, France.**

Stéphane Potteaux

**Inserm, Délégation régionale Paris Île-de-France Centre Nord, Paris, France.**

Stéphane Potteaux

**Université Bourgogne-Europe, INSERM, CTM UMR1231, DesCarTes, 21000 Dijon, France.**

Olivier Micheau & Abdelmnim Radoua

## Contributions

MMC – Conceptualization, methodology, investigation, data analysis, and writing, reviewing and editing; LC- methodology, data analysis, reviewing; AA- conceptualization, methodology, data analysis, writing, reviewing and editing; AB- methodology, data analysis and reviewing; AB-methodology, data analysis and reviewing; JC- methodology and data analysis; CJ- reviewing and editing; AAR- methodology, data analysis and reviewing; HE- methodology; JWD- methodology, data analysis and reviewing; CH- methodology; LC- methodology; KS- reviewing; AR-methodology; CM- methodology; EB- methodology; CBR- methodology, data analysis and reviewing; MBH- reviewing; YM- reviewing; SP- methodology, data analysis and reviewing; OM-methodology, data analysis and reviewing; SD- reviewing; JD- supervision, conceptualization, methodology, investigation, data analysis, and writing, reviewing and editing; JTD-JD- supervision, conceptualization, methodology, investigation, data analysis, and writing, reviewing and editing.

## Corresponding authors

Correspondence to Jérôme Devy

## Ethics declarations

### Ethics approval and consent to participate

All patients provided informed oral consent and signed a non-opposition form and the study was approved by a local ethics committee (CRB-Institut Godinot, approval 170551/1371F). All animal experimental procedures were conducted after obtaining authorization from the French

Ministry of the Research (authorization #8140v7) and at an accredited scientific research animal facility (E-51-454-2). All experience are done on accordance with rules of local ethical committee (CEEA-51).

