## Supplementary_data for "LRP-1 promotes tumor progression of triple negative breast cancers by coordinating extracellular matrix remodeling and immune cell infiltration"

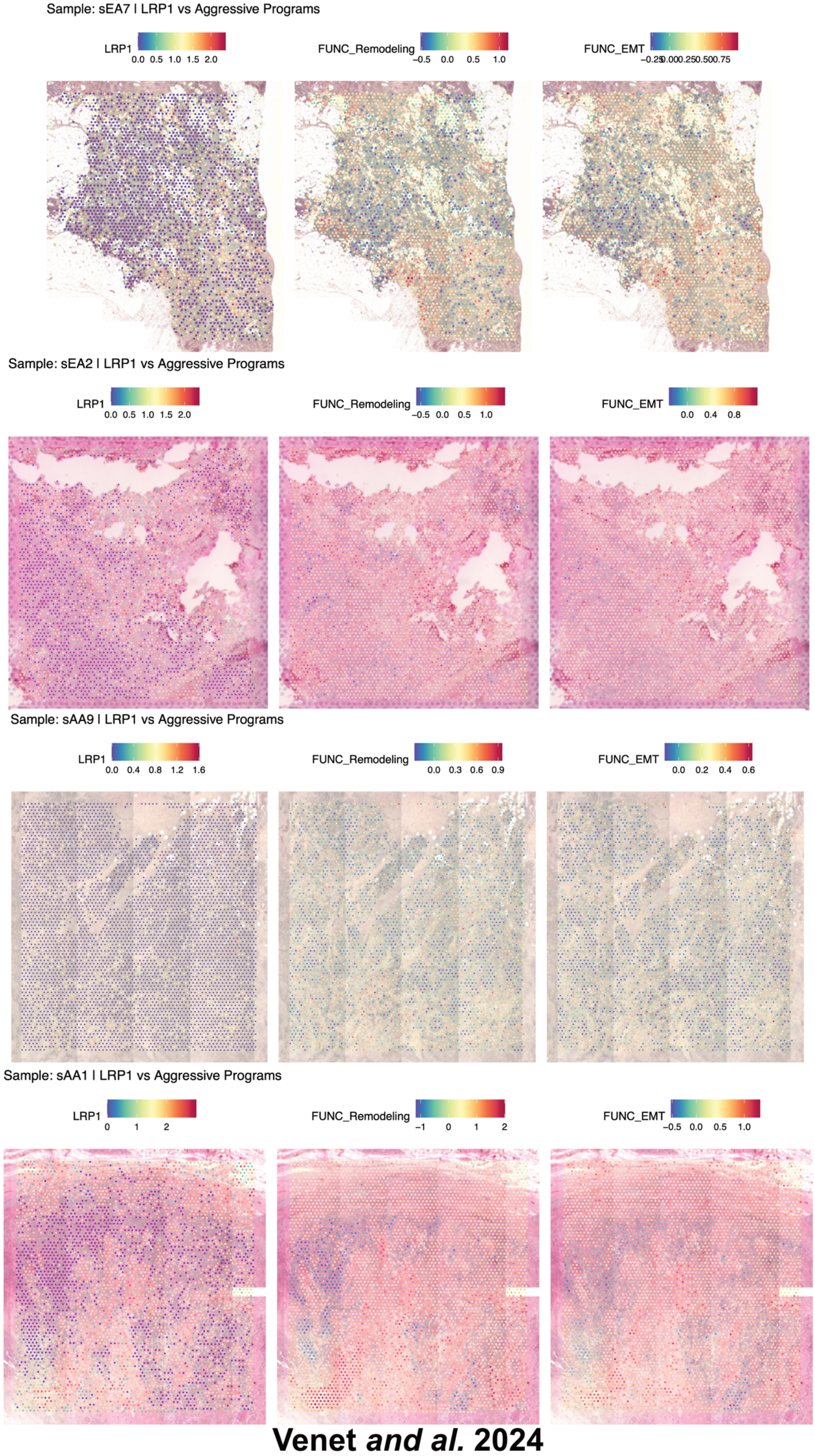


<u>Additional  Figure 1</u>: **Venet and al spatial transcriptomics on TNBC and LRP-1 : all samples**

Spatial transcriptomics from Venet *and al.* (2024) across nine TNBC samples reveals non-uniform, regionally concentrated LRP-1 expression, enriched at infiltrating tumor fronts and stromal-adjacent niches.

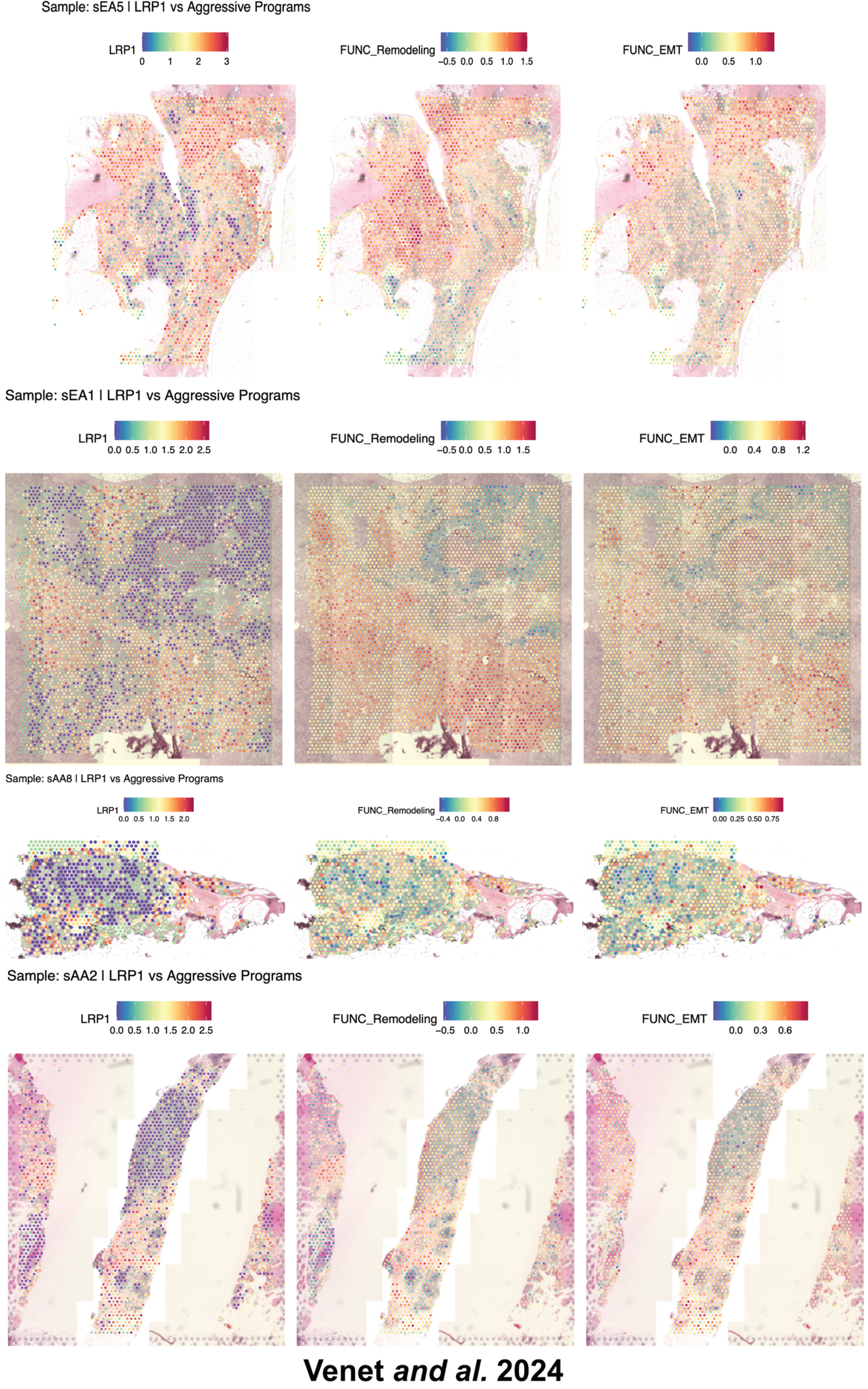


Additional Figure 2: **Venet et al spatial transcriptomics on TNBC and LRP-1 : all samples (continued)**

Spatial transcriptomics from Venet *and al.* (2024) across nine TNBC samples reveals non-uniform, regionally concentrated LRP-1 expression, enriched at infiltrating tumor fronts and stromal-adjacent niches (continued).

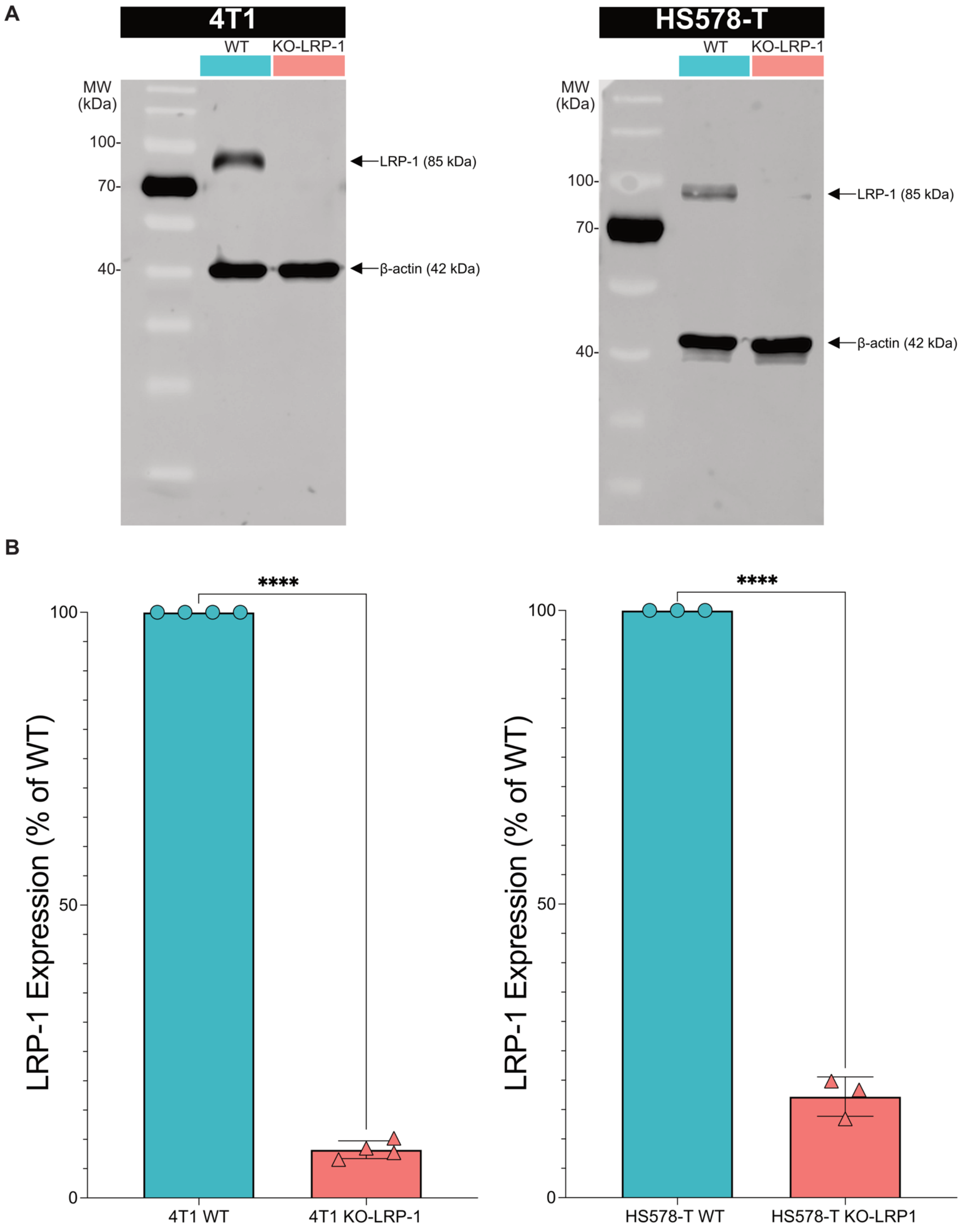


Additional Figure 3: **Validation of KO-LRP-1 models.**

**A:** Representative Western blot images showing LRP-1 protein expression in 4T1 WT/KO-LRP-1 cells (left panel) and in HS578-T WT/KO-LRP-1 cells (right panel), with β-actin serving as an endogenous control. **B:** Densitometric analysis of Western blots showing the relative change in protein expression observed in 4T1 LRP-1 KO cells compared to 4T1 WT cells (left panel) and in HS578-T LRP-1 KO cells compared to HS578-T WT cells (right panel). Data are shown as bar graphs and presented as mean ± SD based on data from three (HS578-T WT/KO-LRP-1) or four (4T1 WT/KO-LRP-1) independent experiments. Statistical analysis was performed using an unpaired two-tailed Student's *t*-test.

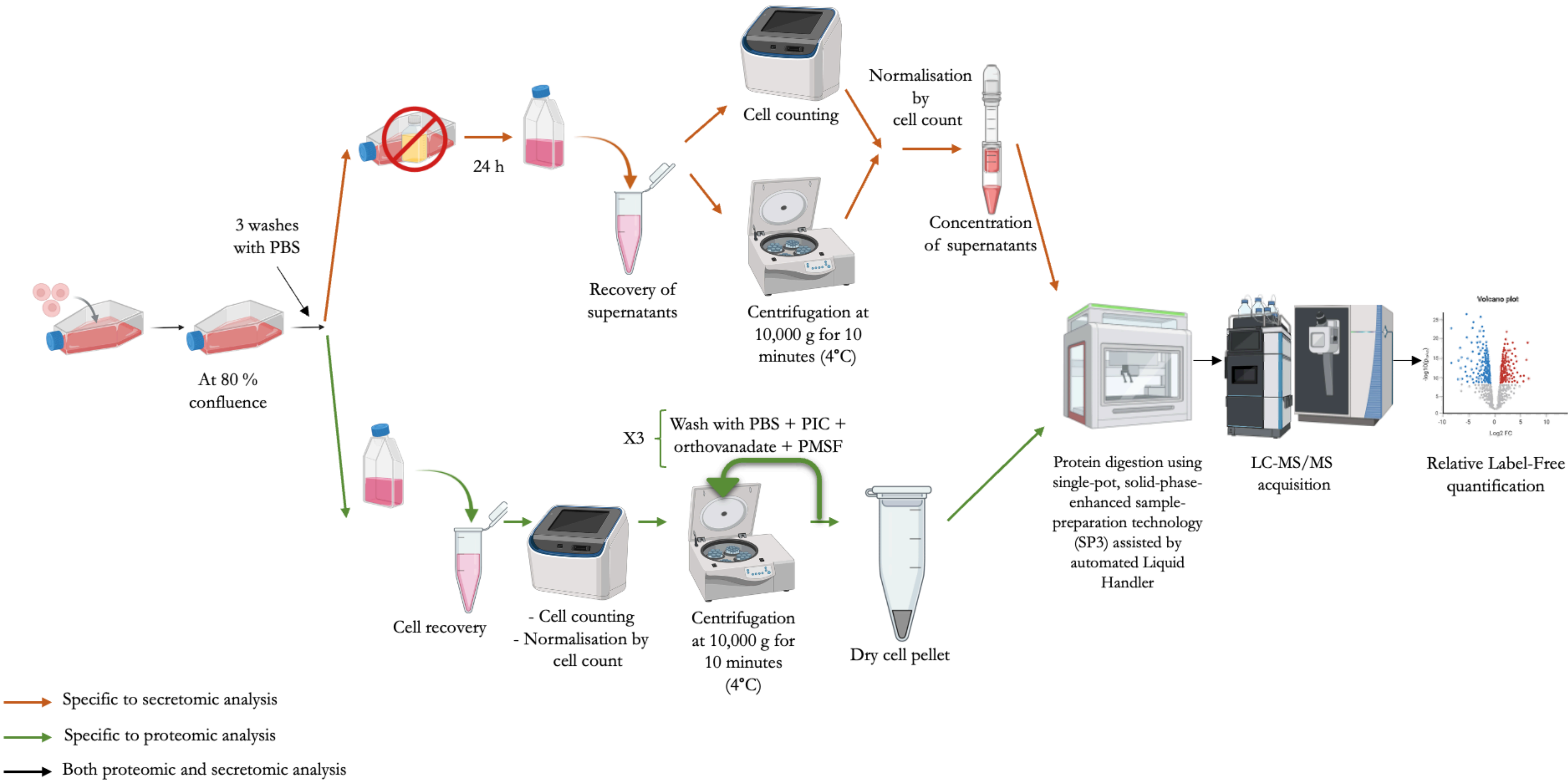


Additional Figure 4: **Sample preparation protocol for proteomic and secretomic analyses.**

The orange arrows indicate steps specific to sample preparation for secretome analyses, the green arrows indicate steps specific to proteome analyses, and the black arrows indicate common steps. The supernatants for secretome analyses were concentrated using Vivaspin columns with a cut-off of 10 Kda.

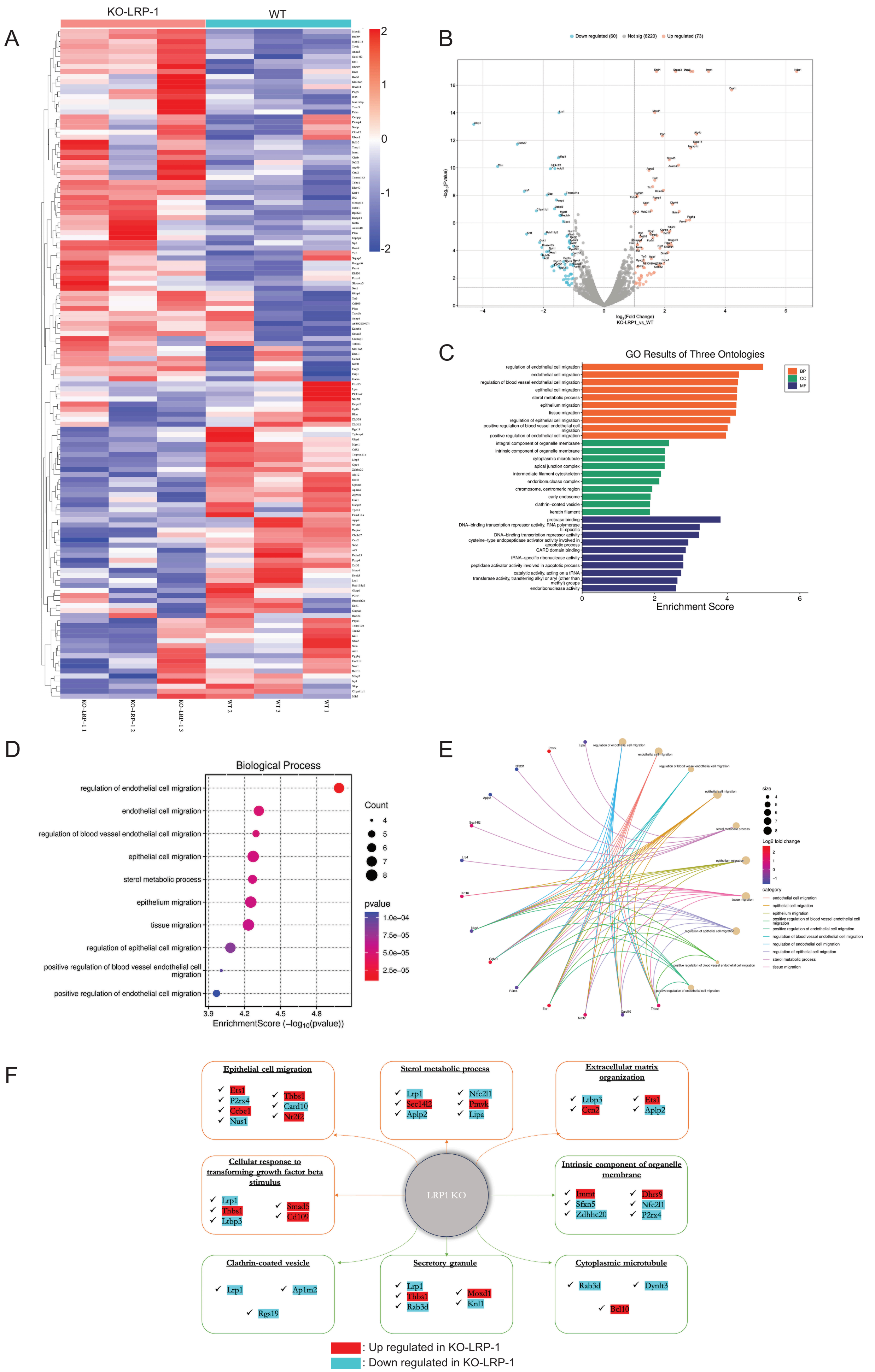
A
KO-LRP-1
WT
2
1
0
-1
-2
KO-LRP-1 1
KO-LRP-1 2
KO-LRP-1 3
WT 2
WT 3
WT 1
B
Down regulated (60)
Not sig (6220)
Up regulated (73)
-log10(Pvalue)
log2(Fold Change)
KO-LRP1_vs_WT
C
GO Results of Three Ontologies
BP
CC
MF
regulation of endothelial cell migration
endothelial cell migration
regulation of blood vessel endothelial cell migration
epithelial cell migration
sterol metabolic process
epithelium migration
tissue migration
regulation of epithelial cell migration
positive regulation of blood vessel endothelial cell migration
positive regulation of endothelial cell migration
integral component of organelle membrane
intrinsic component of organelle membrane
cytoplasmic microtubule
apical junction complex
intermediate filament cytoskeleton
endoribonuclease complex
chromosome, centromeric region
early endosome
clathrin-coated vesicle
keratin filament
protease binding
DNA-binding transcription repressor activity, RNA polymerase II-specific
DNA-binding transcription repressor activity
cysteine-type endopeptidase activator activity involved in apoptotic process
CARD domain binding
tRNA-specific ribonuclease activity
peptidase activator activity involved in apoptotic process
catalytic activity, acting on a tRNA
transferase activity, transferring alkyl or aryl (other than methyl) groups
endoribonuclease activity
Enrichment Score
D
Biological Process
regulation of endothelial cell migration
endothelial cell migration
regulation of blood vessel endothelial cell migration
epithelial cell migration
sterol metabolic process
epithelium migration
tissue migration
regulation of epithelial cell migration
positive regulation of blood vessel endothelial cell migration
positive regulation of endothelial cell migration
Count
4
5
6
7
8
pvalue
1.0e-04
7.5e-05
5.0e-05
2.5e-05
3.9
4.2
4.5
4.8
EnrichmentScore (−log10(pvalue))
E
size
Log2 fold change
category
endothelial cell migration
epithelial cell migration
epithelium migration
positive regulation of blood vessel endothelial cell migration
positive regulation of endothelial cell migration
regulation of blood vessel endothelial cell migration
regulation of endothelial cell migration
regulation of epithelial cell migration
sterol metabolic process
tissue migration
F
Epithelial cell migration
Ets1
P2rx4
Ccbe1
Nus1
Thbs1
Card10
Nr2f2
Sterol metabolic process
Lrp1
Sec14l2
Aplp2
Nfe2l1
Pmvk
Lipa
Extracellular matrix organization
Ltbp3
Ccn2
Ets1
Aplp2
Cellular response to transforming growth factor beta stimulus
Lrp1
Thbs1
Ltbp3
Smad5
Cd109
LRP1 KO
Intrinsic component of organelle membrane
Immt
Sfxn5
Zdhhc20
Dhrs9
Nfe2l1
P2rx4
Clathrin-coated vesicle
Lrp1
Ap1m2
Rgs19
Secretory granule
Lrp1
Thbs1
Rab3d
Moxd1
Knl1
Cytoplasmic microtubule
Rab3d
Dynlt3
Bcl10
: Up regulated in KO-LRP-1
: Down regulated in KO-LRP-1

Additional Figure 5: **LRP-1 drives proteomic rewiring and pathway-level changes.**

Overview of panels showing **A:** Heatmap of 133 proteins differentially expressed between WT and LRP-1 KO cells. Values are logarithmic scale of fold change from -2 to 2, blue = decreased, red = increased. **B:** Volcano plot of differential protein expression between WT and LRP-1 KO cells X = $\log_2$ fold change, Y = $-\log_{10}$ p-value; 60 proteins significantly down-regulated and 73 significantly up-regulated in KO (significant points colored). **C:** Bar plot of S-scores summarizing Gene Ontology enrichment across three groups : biological processes (BP, orange), cellular components (CC, green) and molecular functions (MF, blue). Each bar represents the S-score for an individual enriched term. **D:** Dot plot of S-scores for enriched biological processes. Point color indicates the p-value and point size indicates the number of LRP-1–regulated proteins that contribute to each term. **E:** Cnetplot linking enriched biological processes (BP) to LRP-1–regulated proteins. Term nodes are grouped by BP categories and sized proportionally to the number of LRP-1–regulated proteins mapping to each term (larger = more proteins). Gene nodes (proteins) are colored by $\log_2$ fold change (range $-1 \rightarrow +2$) (blue = lower abundance in KO; red = higher abundance in KO). Only terms and proteins meeting the significance thresholds are shown. **F:** Gene Set Enrichment Analysis and the pathways most affected by LRP-1 loss in the proteome, each annotated with its contributing modulated proteins and a color scale indicating relative abundance (blue = decreased in LRP-1 KO; red = increased in LRP-1 KO.

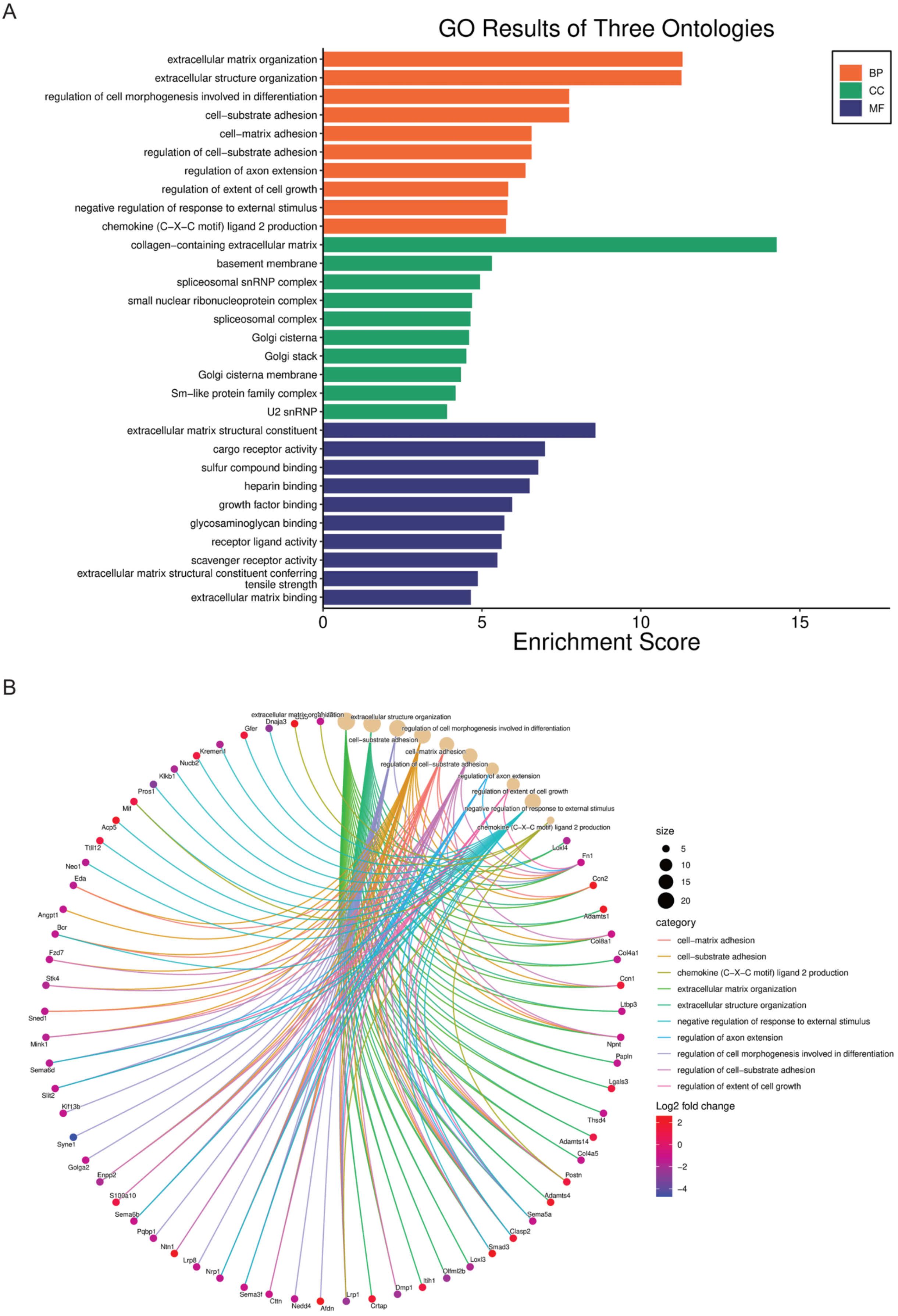


Additional Figure 6: **LRP-1 affects various secretome pathways.**

**A:** Bar plot of S-scores summarizing Gene Ontology enrichment across three groups : biological processes (BP, orange), cellular components (CC, green) and molecular functions (MF, blue). Each bar represents the S-score for an individual enriched term. **B:** Cnetplot linking enriched biological processes (BP) to LRP-1 regulated proteins. Term nodes are grouped by BP category and sized proportionally to the number of LRP-1 regulated proteins mapping to each term (larger = more proteins). Gene nodes (proteins) are colored by $\log_2$ fold change (range -4 → +2) (blue = lower abundance in KO; red = higher abundance in KO). Only terms and proteins meeting the significance thresholds are shown.

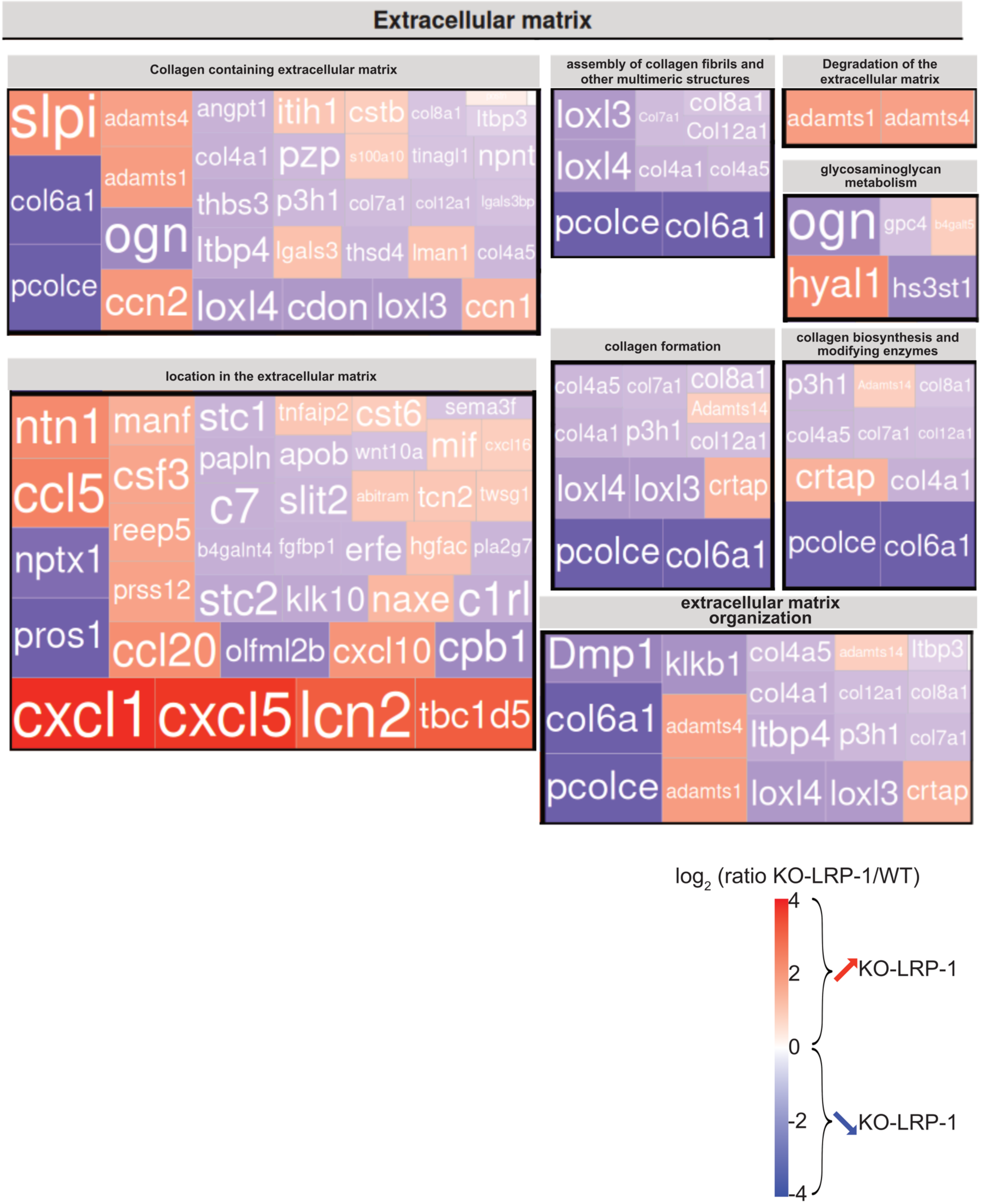


Additional Figure 7: **Subdivision of ECM processes affected by LRP-1.**

Treemap of eight ECM-related sub-pathways most impacted by LRP-1 deletion. Each rectangle represents one LRP-1–regulated protein mapped to the corresponding sub-pathway; rectangles are grouped by sub-pathway. Color indicates the protein $log_2$(KO/WT) ratio (scale −4 → +4; blue = decreased in KO, red = increased in KO). Only proteins and terms meeting the significance criteria are shown (GSEA across Hallmark, GOCC and IPA).

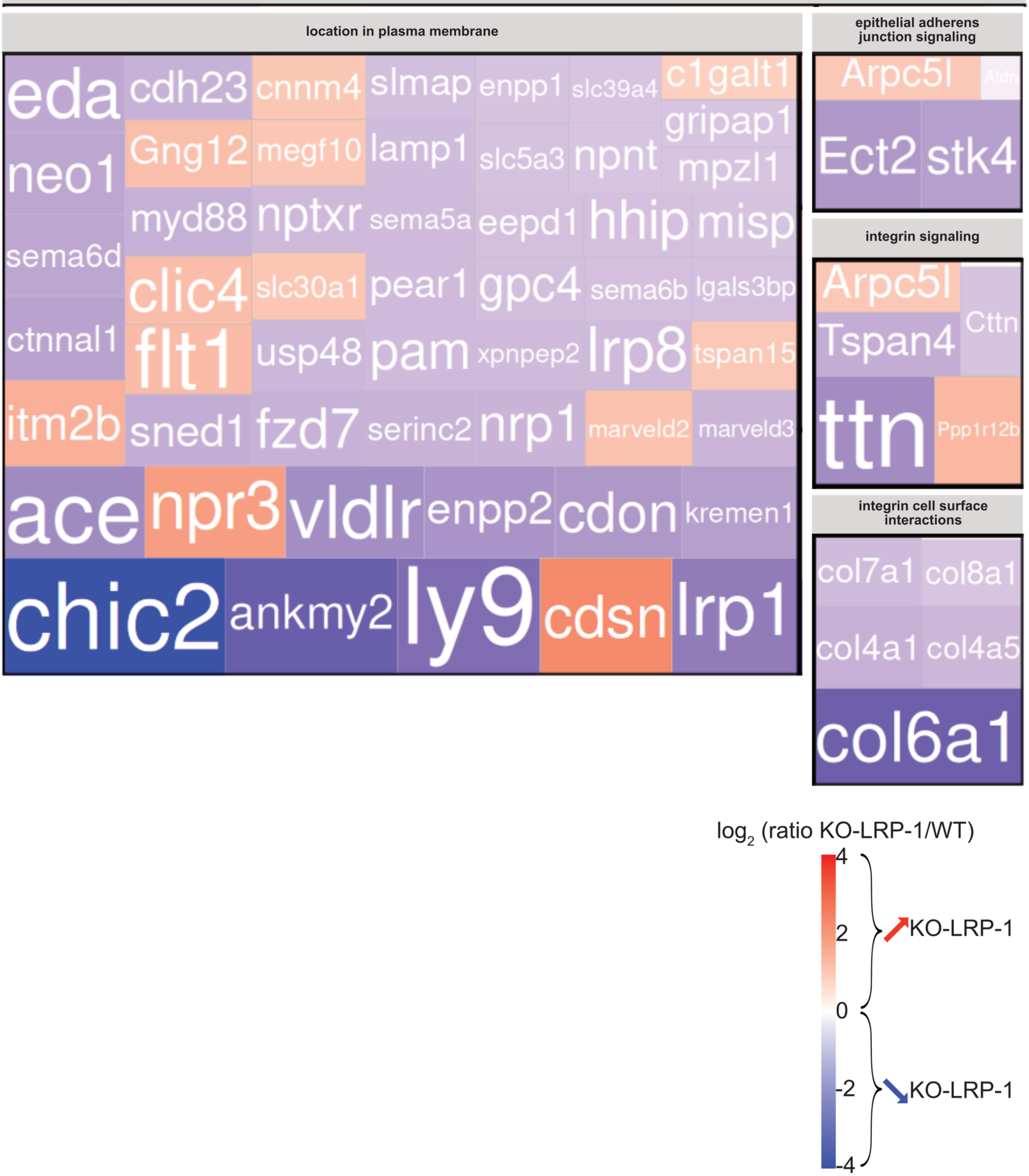


<u>Additional Figure 8:</u> **Subdivision of plasma membrane processes affected by LRP-1.**

Treemap of four plasma membrane related sub-pathways most impacted by LRP-1 deletion. Each rectangle represents one LRP-1–regulated protein mapped to the corresponding sub-pathway; rectangles are grouped by sub-pathway. Color indicates the protein $\log_2$(KO/WT) ratio (scale −4 → +4; blue = decreased in KO, red = increased in KO). Only proteins and terms meeting the significance criteria are shown (GSEA across Hallmark, GOCC and IPA).

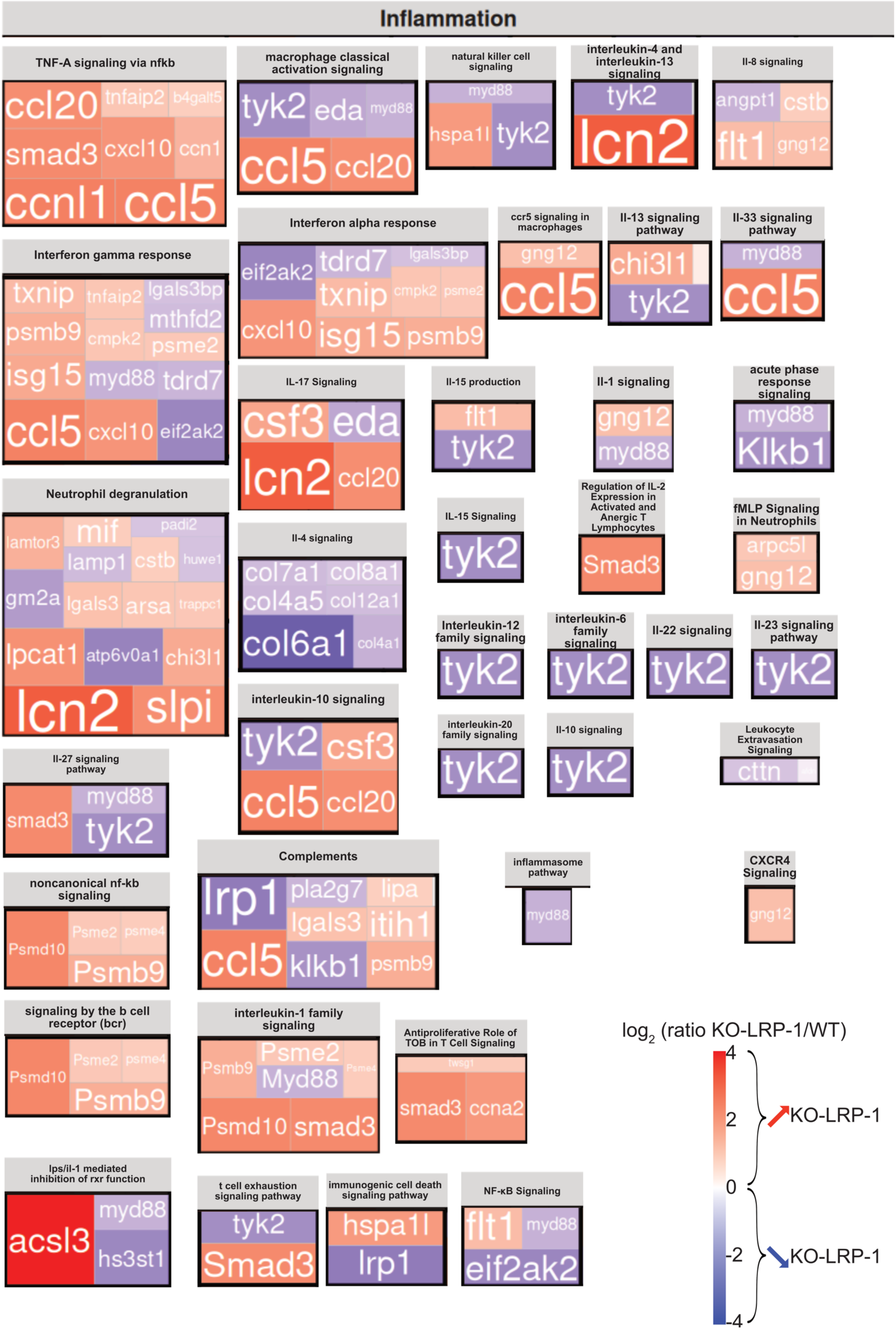
Inflammation
TNF-A signaling via nfkb
ccl20 tnfaip2 b4galt5
smad3 cxcl10 ccn1
ccnl1 ccl5
macrophage classical activation signaling
tyk2 eda myd88
ccl5 ccl20
natural killer cell signaling
myd88
hspa1l tyk2
interleukin-4 and interleukin-13 signaling
tyk2
lcn2
Il-8 signaling
angpt1 cstb
flt1 gng12
Interferon gamma response
txnip tnfaip2 lgals3bp
psmb9 cmpk2 mthfd2 psme2
isg15 myd88 tdrd7
ccl5 cxcl10 eif2ak2
Interferon alpha response
eif2ak2 tdrd7 lgals3bp
txnip cmpk2 psme2
cxcl10 isg15 psmb9
ccr5 signaling in macrophages
gng12
ccl5
Il-13 signaling pathway
chi3l1
tyk2
Il-33 signaling pathway
myd88
ccl5
IL-17 Signaling
csf3 eda
lcn2 ccl20
Il-15 production
flt1
tyk2
Il-1 signaling
gng12
myd88
acute phase response signaling
myd88
Klkb1
Neutrophil degranulation
lamtor3 mif padi2
lamp1 cstb huwe1
gm2a lgals3 arsa trappc1
lpcat1 atp6v0a1 chi3l1
lcn2 slpi
Il-4 signaling
col7a1 col8a1
col4a5 col12a1
col6a1 col4a1
IL-15 Signaling
tyk2
Regulation of IL-2 Expression in Activated and Anergic T Lymphocytes
Smad3
fMLP Signaling in Neutrophils
arpc5l
gng12
Interleukin-12 family signaling
tyk2
interleukin-6 family signaling
tyk2
Il-22 signaling
tyk2
Il-23 signaling pathway
tyk2
interleukin-10 signaling
tyk2 csf3
ccl5 ccl20
interleukin-20 family signaling
tyk2
Il-10 signaling
tyk2
Leukocyte Extravasation Signaling
cttn
Il-27 signaling pathway
smad3 myd88
tyk2
Complements
lrp1 pla2g7 lipa
lgals3 itih1
ccl5 klkb1 psmb9
inflammasome pathway
myd88
CXCR4 Signaling
gng12
noncanonical nf-kb signaling
Psmd10 Psme2 psme4
Psmb9
signaling by the b cell receptor (bcr)
Psmd10 Psme2 psme4
Psmb9
interleukin-1 family signaling
Psmb9 Psme2 Psme4
Myd88
Psmd10 smad3
Antiproliferative Role of TOB in T Cell Signaling
twsg1
smad3 ccna2
log2 (ratio KO-LRP-1/WT)
4
2
0
-2
-4
KO-LRP-1
KO-LRP-1
lps/il-1 mediated inhibition of rxr function
acsl3 myd88
hs3st1
t cell exhaustion signaling pathway
tyk2
Smad3
immunogenic cell death signaling pathway
hspa1l
lrp1
NF-κB Signaling
flt1 myd88
eif2ak2

Additional Figure 9: **Subdivision of inflammation processes affected by LRP-1.**

Treemap of thirty-nine inflammation related sub-pathways most impacted by LRP-1 deletion. Each rectangle represents one LRP-1–regulated protein mapped to the corresponding sub-pathway; rectangles are grouped by sub-pathway. Color indicates the protein $\log_2$(KO/WT) ratio (scale $-4 \rightarrow +4$; blue = decreased in KO, red = increased in KO). Only proteins and terms meeting the significance criteria are shown (GSEA across Hallmark, GOCC and IPA).

### Additional Materiels 1

#### Immunohistochemistry on patient biopsy and on mouse tumors

Paraffin-embedded tissues were obtained from TNBC female patients. All patients provided informed oral consent and signed a non-opposition form and the study was approved by a local ethics committee (CRB-Institut Godinot, approval 170551/1371F). Mouse tumors were fixed 48 hours in 4% paraformaldehyde solution and then embedded in paraffin to form blocks. Tissues were cut into 5 µm sections using Microm HM 335 E Microtome (MM France). Tissues were deparaffinized and rehydrated using xylene and a graded series of ethanol. Antigen retrieval was performed in ImmunoRetriever with citrate (pH=6) (Bio SB ImmunoRetriever 20X with citrate) and with TintoRetriever pressure cooker (Bio SB) for 20 minutes. Permeabilization was obtained by a 10-minute incubation with 0.1% Triton X-100. Mouse and Rabbit Specific HRP/DAB IHC detection kit, Micro-polymer (Abcam) was used according to the manufacturer's protocol. Briefly, endogenous peroxidase activity was blocked by 10 minutes incubation with Hydrogen Peroxide reagent followed by incubation with Protein Block reagent. Samples were then incubated overnight with a suitable primary antibody (see Additionnal_table_1 for more details) at 4°C. Tissues were further treated with the secondary antibody using HRP-conjugated DAB substrate, counterstained in hematoxylin for 30 seconds and mounted in BioMount DPX New (Biognost). The quantification of DAB staining was performed with Qpath which can detect and measure the stained pixels. Data are presented as a table (for patient biopsy) and plot as a bar graph with mean ± SD (for mouse tumors). Statistical comparisons were performed using a two-tailed Student's t-test : $P < 0.05$ (*).

#### Public TNBC dataset acquisition and pre-processing

Three independent, publicly available transcriptomic datasets were interrogated to characterise *LRP-1* expression across breast cancer subtypes and within the TNBC tumor microenvironment (TME).

##### Single-cell RNA sequencing:

**Wu *and al.* (2021).** The breast cancer single-cell atlas generated by Wu *and al.* (Nature Genetics, 2021; GSE176078) was retrieved from the Gene Expression Omnibus (GEO). This dataset comprises 24,271 single cells profiled by 10x Genomics Chromium from primary human breast tumors, including TNBC. Cell-type annotations provided by the original authors were retained and encompassed 18 populations: B cells, $CD4^+$ and $CD8^+$ T cells, dendritic plasmacytoid-like cells (dPVL), endothelial cells, epithelial basal, epithelial basal cycling, epithelial luminal mature, inflammatory cancer-associated fibroblasts (iCAFs), myoCAFs, myeloid cells, myoepithelial cells, NK cells, NKT cells, plasma cells, T cells cycling, T cells unassigned, Tfh cells, and regulatory T cells (T-regs). Dimensionality reduction and visualisation were performed using t-distributed stochastic neighbour embedding (tSNE), as originally computed by the dataset authors. *LRP-1* mRNA expression values ($log_2$-fold) were overlaid onto the tSNE embedding to identify cell populations with preferential expression.

**Chung *and al.* (2017).** The breast cancer single-cell transcriptomic dataset generated by Chung *and al.* (Nature Communications, 2017; GSE75688) was retrieved from GEO. This dataset discriminates four breast cancer molecular subtypes: $ER^+/HER2^+$, $ER^+$, $HER2^+$, and TNBC, enabling inter-subtype comparisons. *LRP-1* mRNA expression levels (counts per million, CPM) were overlaid onto the tSNE projection to assess subtype-specific enrichment.

**Bulk transcriptomics:**

**DLDCC (2022)[1].** The DLDCC 2022 breast cancer dataset, accessible via cBioPortal for Cancer Genomics (*https://www.cbioportal.org*), was used for bulk transcriptomic analysis stratified by TNBC molecular subtype. *LRP-1* mRNA expression was compared across the six Lehmann TNBC molecular subtypes — basal-like 1 (BL1), basal-like 2 (BL2), immunomodulatory (IM), mesenchymal (M), mesenchymal stem-like (MSL), and luminal androgen receptor (LAR), as assigned by the dataset authors. Mutation type annotations and copy number variation (CNV) data available within the dataset were incorporated for per patient visualisation.

**Spatial transcriptomic data acquisition and preprocessing**

Spatial transcriptomic (ST) data from nine TNBC tumor tissue sections were obtained from the publicly available dataset generated by Venet *and al.[2]* (2024; hereafter Venet *and al.* TNBC ST dataset), accessible via Zenodo (https://zenodo.org). Samples were acquired using the 10x Genomics Visium platform, providing spatially resolved gene expression data co-registered with haematoxylin and eosin (H&E) histology images. Data processing was performed in R (v4.3.2) using the Seurat package (v5.0.3).

Raw feature-barcode matrix data were loaded per sample using *Load10X_Spatial()*. Quality control (QC) filtering was applied to remove low-quality spots: spots with fewer than 200 detected genes or with mitochondrial transcript content exceeding 20% were excluded. Normalisation was performed using the regularised negative binomial regression method implemented in *SCTransform()* (Hafemeister & Satija, Genome Biology, 2019)[3], applied to the Spatial assay with default parameters (variance-stabilising transformation, regression of sequencing depth). Principal component analysis (PCA) was subsequently applied using the 3,000 most variable features identified by SCTransform, retaining the top 30 principal components (PCs) for downstream analyses. Uniform Manifold Approximation and Projection (UMAP) was computed from the PCA embedding (dims = 1–30). Unsupervised spot clustering was performed using a shared-nearest-neighbour (SNN) graph (*FindNeighbors()*, dims = 1–30) followed by the Louvain algorithm at resolution 0.5 (*FindClusters()*).

#### Identification of spatially variable features

Spatially variable gene expression features were identified per sample using Moran's I statistic, implemented in *FindSpatiallyVariableFeatures()* (selection.method = 'moransi'; Seurat v5). The top 1,000 variable features identified by SCTransform were ranked by spatial autocorrelation. Moran's I quantifies the degree to which gene expression is spatially clustered relative to a random distribution, providing an unbiased assessment of spatial patterning. Results were used to contextualise the spatial distribution of *LRP-1* and to inform the selection of spatially informative signatures for downstream correlation analyses.

#### Gene signature scoring

Spot-level gene signature scores were computed using the *AddModuleScore()* function in Seurat, which calculates the average expression of a defined gene set relative to a set of randomly sampled background genes (Tirosh *and al.*, Science, 2016)[4]. For each sample, only genes present in the feature matrix were included in scoring (intersect with rownames). A minimum of three detected genes per signature was required for scoring; signatures failing this threshold were excluded. Background gene bins (nbin = 24) and control gene sets (ctrl = 100) were used as implemented by default.

Four categories of gene signatures were assessed:

**TNBC molecular subtypes (Lehmann classification).** Subtype-defining gene sets for BL1, BL2, immunomodulatory (IM), mesenchymal (M), mesenchymal stem-like (MSL), and luminal androgen receptor (LAR) subtypes were derived from Lehmann *and al.* (Journal of Clinical Investigation, 2011)[5] and implemented as curated gene sets (BL1: *MKI67, TOP2A, AURKA, AURKB, BUB1, CDK1, CCNB1, CCNB2, PCNA, BRCA1*; M: *VIM, FN1, COL1A1, SNAI2, TWIST1, ZEB1, ZEB2, CDH2, MMP2, MMP14, LOXL2, TGFB1, POSTN, ACTA2, FAP, PDGFRB*; LAR: *AR, FOXA1, KRT18, KRT19, XBP1, GATA3, AGR2, TFF1, MLPH, NAT1*; and corresponding sets for BL2, IM, and MSL).

**Tumor microenvironment cell populations.** Microenvironmental signatures comprised: cancer-associated fibroblast (CAF: *ACTA2, FAP, COL1A1, COL1A2, COL3A1, PDGFRA, PDGFRB, THY1, POSTN, SPARC, VCAN, FN1, TNC, CXCL12*), stromal (*DCN, LUM, COL1A1, COL1A2, COL3A1, BGN, VCAN, FN1, SPARC*), immune infiltration (*PTPRC, CD3D, CD3E, MS4A1, CD79A, CD74, HLA-DRA, HLA-DRB1, CXCL9, CXCL10, GZMB, NKG7, CCL5*), and vascular (*PECAM1, VWF, KDR, CDH5, FLT1, ENG, CLEC14A*).

**Functional transcriptional programmes.** Functional signature gene sets were curated from MSigDB Hallmark gene sets (Liberzon *and al.*, Cell Systems, 2015)[6] and published literature: epithelial-to-mesenchymal transition (EMT; HALLMARK_EPITHELIAL_MESENCHYMAL_TRANSITION), extracellular matrix (ECM) remodelling (*MMP2, MMP9, MMP11, MMP14, PLAU, PLAUR, CTSB, CTSL, LOX, LOXL1, LOXL2, POSTN, FN1, COL1A1, COL1A2, VCAN, SPARC, THBS1, TIMP1, TIMP2, SERPINE1, ADAMTS1, HAS2, FAP*), hypoxia (HALLMARK_HYPOXIA), inflammatory response (HALLMARK_INFLAMMATORY_RESPONSE), and proliferation (HALLMARK_E2F_TARGETS and HALLMARK_G2M_CHECKPOINT combined). Critically, *LRP-1* was excluded from all signature gene sets to ensure that correlations reflect independent biological programmes rather than circular scoring.

<u>**Spatial correlation and statistical analysis**</u>

Spot-level Spearman rank correlations were computed between *LRP-1* normalised expression (SCTransform-corrected counts) and each gene signature score across all informative spots per sample. Spearman's rho (ϱ) was selected for its robustness to non-normality and outliers inherent to spatial transcriptomic data. Correlations were computed only for spots with $\geq 10$ valid (finite, non-missing) paired observations. Unadjusted two-sided p-values were corrected for multiple testing using the Benjamini–Hochberg (BH) false discovery rate (FDR) procedure applied within each analytical category (TNBC subtypes, microenvironment, functional programmes) per sample. Associations with FDR-adjusted $p < 0.05$ were considered statistically significant. Per-sample Spearman ϱ values were assembled into a signature × sample matrix and visualised as a colour-scaled heatmap (blue: negative association; red: positive association; white: no association). Summary statistics across the nine-sample cohort (mean ϱ, median ϱ, standard deviation, proportion of samples with positive rho, and proportion reaching FDR $< 0.05$) were computed to assess cross-sample consistency of each LRP-1–programme association.

To account for spot-level pseudo-replication arising from spatial autocorrelation and to model inter-sample heterogeneity, a linear mixed-effects model was fitted to the pooled spot-level dataset using the lme4 package (v1.1-35; Bates *and al.*, Journal of Statistical Software, 2015)[7]: *LRP-1 ~ EMT_Score + ECM_Remodeling_Score + CAF_Score + (1|sample)*, where sample was included as a random intercept to account for between-sample variation. Model fixed-effect estimates and 95% confidence intervals are reported.

**<u>Visualisation</u>**

Spatial gene expression maps were generated using *SpatialFeaturePlot()* (Seurat v5; pt.size.factor = 1.6) overlaid on matched H&E histology images. LRP-1 expression was visualised using a continuous colour gradient (grey: low expression; red: high expression). Signature score spatial maps were rendered with programme-specific colour gradients to facilitate visual differentiation. Scatter plots of spot-level *LRP-1* expression versus signature scores were produced using ggplot2 (v3.5.0), with a linear regression trend line overlaid; Spearman $\rho$ and FDR-adjusted p-values were annotated on each plot. Cross-sample heatmaps of Spearman $\rho$ values were generated using ggplot2 with a diverging colour scale centred at zero (blue–white–red, limits $\pm 1$). Consistency dot plots displaying per-sample $\rho$ values across signatures were generated using ggplot2 with faceting by analytical category. All figures were assembled and annotated using the patchwork package (v1.2.0) and finalised in Adobe Illustrator.

**<u>Software and reproducibility</u>**

All bioinformatic analyses were performed in R (v4.3.2) with the following core packages: Seurat v5.0.3, ggplot2 v3.5.0, dplyr v1.1.4, patchwork v1.2.0, tidyr v1.3.0, purrr v1.0.2, lme4 v1.1-35, pheatmap v1.0.12, msigdbr v7.5.1, Rfast2 v0.1.5, and ape v5.7-1. Random seeds were set to 123 for all stochastic steps (SCTransform normalisation, UMAP embedding, module scoring). All analysis code and intermediate R objects will be deposited in a public repository upon acceptance of this manuscript.

#### Cell culture and LRP-1 knockout model generation

We obtained the murine mammary carcinoma cell line 4T1 (ATCC®, catalog : CRL-2539™) and the human breast cancer cell line HS578-T (ATCC®, catalog : HTB-126™) from the American Type Culture Collection (ATCC, Manassas, VA, USA) and maintained them following the supplier's recommendations. We generated LRP-1 knockout models, without clonal selection, of the 4T1 and HS578-T cell lines using Crispr/Cas9 strategy as described in our previous study[8]. The models were revalidated once again in this study using Western blots (fig_additional_3).

#### Mass spectrometry-based label-free quantitative proteomics and secretomics

The preparation of samples, of three independent experiments, for proteomic and secretomic analyses is described in fig_additional_4. After extraction from the pellets (proteomes) and in the secretomes, proteins were precipitated by adding trichloroacetic acid (TCA) to a final concentration of 10% (w/v), followed by incubation on ice for 30 minutes. Samples were then centrifuged at 15,000 × g for 30 minutes at 4°C. The resulting protein pellets were washed once with cold acetone and centrifuged again at 15,000 × g for 15 minutes at 4°C. After removal of the supernatant, pellets were air-dried briefly and resuspended in 150 mM Tris-HCl buffer (pH 8.5) containing 1% SDS and a protease inhibitor cocktail. Proteins were reduced with 30 mM dithiothreitol (DTT) for 1 hour at 56°C, and subsequently alkylated with 90 mM iodoacetamide (IAA) for 1 hour at room temperature in the dark. Proteins were then desalted and digested by the trypsin overnight 37°C using the Single-pot, Solid-phase-enhanced Sample Preparation (SP3) method. NanoLC-MS/MS analysis were performed using a Vanquish Neo UHPLC System (Thermo Scientific) associated to Orbitrap Exploris™ 480, (see the parameters used [9]). Protein identification and label-free quantification (LFQ) were done in Proteome Discoverer 3.2. The CHIMERYS node using the prediction model inferys_4.7.0 fragmentation was used to identify proteins in batch mode by searching against a UniProt *Mus musculus* protein database (54750 entries, released August 2025). (See the parameters used [9]). Quantitative data were considered for master proteins, quantified by a minimum of 2 unique peptides and normalize following total peptide quantity normalization. Protein identifications were retained at an FDR of <1%, and missing values were imputed using the Low Abundance Resampling approach. Differential abundance was defined by $P < 0.05$ and a fold change of ≤0.5 or ≥2.0. The mass spectrometry proteomics data have been deposited to the ProteomeXchange Consortium via the PRIDE [10] partner repository with the dataset identifiers: PXD072836 (Total proteome) and PXD072843 (Secretomes). All figures presented were created using bioinformatics analysis tools such as SRplot [11] or Ingenuity Pathways.

#### 2D Migration assay

We assessed collective cell migration using silicone migration inserts (catalog : 80206; Culture-Insert 2 Well in μ-Dish 35 mm, low, IBIDI) following the manufacturer's instructions. 4T1 WT and 4T1 LRP-1-knockout cells were seeded at $6.5.10^4$ cells per well in migration-insert dishes in RPMI-1640 (catalog : 7240047, Thermo Fisher Scientific, Waltham, MA, USA) supplemented with 10% fetal bovine serum (FBS) (catalog : F7524-500ml, batch #0001672583, Sigma-Aldrich). Human HS578-T WT and HS578-T KO-LRP-1 cells were seeded at $8.10^4$ cells in high-glucose DMEM (4.5 g/L) (catalog : 31966021, Thermo Fisher Scientific, Waltham, MA, USA) supplemented with 10% FBS and 0.01 mg/mL human insulin (catalog : 11508856, Gibco™). Plates were incubated for 24 h at 37 °C in a humidified 5% CO2 incubator to allow attachment and formation of a confluent monolayer. Prior to wound formation, monolayers were washed three times with pre-warmed PBS to remove residual growth medium. To prevent proliferation from confounding wound closure measurements, cells were serum-starved and treated with mitomycin C (cytostatic agent) (catalog : M4287, Merck) for 2 h at 10 μg/mL. Following three additional washes with pre-warmed PBS, silicone inserts were removed using sterile forceps to create a reproducible, cell-free gap. Cells were then incubated in migration medium containing 1% FBS (and 0.01 mg/mL human insulin for HS578-T cells) for the duration of the assay. Phase-contrast images were acquired immediately after insert removal (T = 0 h) and at 24 h (T = 24 h) using an inverted microscope (evos cell imaging systems, Thermo Fisher Scientific, Waltham, MA, USA). Wound areas were measured from images using image-analysis software (ImageJ/Fiji, Wound_healing_size_tools_uptaded) and wound closure was expressed relative to the initial wound area (T = 0 h normalized to 100%). Data are presented as mean ± SD. Statistical comparisons were performed using a two-tailed Student's t-test : $P < 0.0001$ (****).

#### 2D monodisperse migration assay

The migration of monodisperse 4T1 and HS578-T WT and LRP-1 KO cells was assessed 24 hours were spent allowing the cells to migrate after 25,000 cells were seeded into each well of a 12-well plate. Cell tracking was done in ImageJ (NIH) using the Manual Tracker and chemotaxis_tool plugins, Euclidean distance, cumulative distance, and velocity were calculated. Data is shown as scatter plots and trajectories as radar charts. For two-group comparisons, the two-tailed Student's t-test was employed; significance is indicated by $P<0.01$ (**), $P < 0.001$ (***), and $P < 0.0001$ (****).

#### 3D migration in collagen gels

3D collagen gels were produced as described in the study by Le and al.[12] using the appropriate media. For each condition we used 200,000 HS578-T (WT or KO-LRP-1) cells or 150,000 4T1 (WT or KO-LRP-1) cells. plates were returned to the incubator for 24 h prior to imaging. Time-lapse imaging was performed on a ZEISS Axiovert 200 inverted microscope equipped with an environmental chamber to maintain 37 °C and 5% $CO_2$. For each well we recorded three fields of view using a 10X objective; z-stacks of 80 planes were acquired every 20 min for 24 h. Image sequences were analysed with C4D tracking software (Inserm, Reims) : for each position we tracked 12 isolated, viable cells (≈215 cells per condition; ≈860 cells in total across experiments). Cell speed was calculated from frame-to-frame displacements using the formula:

$$Speed = 3\sqrt{\{(dx \times 1.25)^2 + (dy \times 1.25)^2 + (dz \times 10)^2\}}$$

where dx and dy are displacements in x and y in pixels, dz is displacement in z in z-steps, 1.25 µm/pixel and 10 µm/z-step convert image units to micrometres, and the factor 3 converts the displacement measured every 20 min to µm/h. Results were plotted as scatter dot plots, with each dot representing an individual cell and a horizontal line indicating the median migration speed. Statistical comparisons between groups were performed using the nonparametric Mann–Whitney test. Statistical significance is denoted as follows: $P < 0.0001$ (****).

#### Cell invasion assay

We evaluated cellular invasiveness using Matrigel®-coated transwell inserts (8-µm pores; catalog : 662638 Greiner Bio-One) as follows. Matrigel® (catalog : 354230, Corning) was thawed on ice, diluted to 300 µg/mL in ice-cold phosphate-buffered saline (PBS, 4 °C) and left to equilibrate overnight at 4 °C on a rotary mixer. We deposited 100 µL of the diluted Matrigel® onto each polyvinyl membrane and allowed the coating to dry for 24 h. Prior to use, membranes were rehydrated with 100 µL PBS for 30 min. Cells were resuspended in serum-reduced medium (RPMI (4T1) or DMEM (HS578-T) containing 2.5% FCS, v/v, and 0.2% BSA, w/v) and 200 µL of this suspension was placed in the upper chamber (75,000 HS578-T cells or 75,000 4T1 cells per insert). The lower chamber was filled with 800 µL RPMI (4T1) or DMEM (HS578-T) supplemented with 10% FCS (v/v) and 2% BSA (w/v) to act as a chemoattractant. Chambers were incubated for 24 h at 37 °C in a humidified atmosphere with 5% $CO_2$. After incubation, inserts were washed twice with PBS in a Class II biosafety cabinet. Cells were fixed by adding cold methanol (–20 °C) to both compartments (200 µL upper; 800 µL lower), and membranes were washed three times with distilled water. We stained membranes with crystal violet (Sigma-Aldrich, cat. no. HT90132-1L) for 20 min (200 µL upper; 800 µL lower), rinsed repeatedly in distilled water until background staining was removed, and gently removed non-invading cells from the upper surface with a cotton swab. Membranes were excised with a scalpel, transferred to a clear 96-well plate and destained in distilled water containing 10% acetic acid for 2 h. Eluates and blanks (distilled water + 10% acetic acid) were read at 560 nm on a Multiskan FC plate reader (Thermo Fisher Scientific). Absorbance values were blank-corrected before analysis. Unless stated otherwise, each condition was assayed in technical triplicate and the experiments were repeated independently at least three times. When comparing two groups, a two-tailed Student's t-test was used and statistical significance is indicated in the text and figures by asterisks: P < 0.001 (***) and *P* < 0.0001 (****).

#### 3D Spheroid Formation and Analysis

5000 HS578-T and 4T1 cells, (WT and KO), detached with Accutase®, were resuspended in 100 µL of EMEM supplemented with 10% (v/v) FBS and 3.5% (v/v) Matrigel® and seeded into ultra-low attachment 96-well round-bottom plates (Cellstar® Cell-Repellent Surface, Greiner Bio-One, Germany). Plates were then incubated at 37 °C for 3 days to allow spheroid formation (day 0). For spheroid growth assessment, 100 µL of complete culture medium (EMEM + 10% FBS) were added at day 0 to each well, and replaced every 3 days to maintain adequate nutrient supply. Spheroid growth was evaluated at day 14. For spheroid invasion assay, spheroids were generated as described above, except that cells were seeded in 50 µL of medium. At day 0, plates were placed on ice for 15 minutes, and 50 µL of Cultrex™ (Trevigen®, Bio-Techne SAS, France) was added to each well. Following centrifugation (5 min, 300 × g, 4 °C), plates were incubated for 1 h at 37

°C to allow matrix polymerization. Then, 100 μL of complete culture medium were added to each well. Half of the medium was replaced every 3 days, and cell invasion was observed at day 7 using an inverted microscope (EVOS FL Auto 2, Invitrogen, Thermo Fisher Scientific). All analysis were performed using FIJI/ImageJ software. When comparing two groups, a two-tailed Student's t-test was used and statistical significance is indicated in the text and figures by asterisks: P < 0.01 (**), P < 0.001 (***) and *P* < 0.0001 (****).

**HES Coloration and Immunostaining analysis of spheroids**

Spheroids were generated as described above. Twelve spheroids were collected at day 14 and fixed in 4% formaldehyde for 5 min at room temperature and embedded in a cassette using the CytoBlock kit (Fisher Scientific) according to the manufacturer's instructions. Cassette were centrifuged using a Cytospin cytotunnel system at 1,500 rpm, incubated in ethanol overnight, embedded in paraffin, and paraffin blocks were sectioned in 3 μm thickness using a microtome. Hematoxylin–eosin–saffron (HES) colorations were performed in collaboration with the Pathology Department of the Reims University Hospital (CHU de Reims). Immunostaining analysis was performed using an antibody directed against LRP-1 (Merck, HPA004182), E-cadherin, or N-cadherin (see Additionnal_table_1 for more informations).

**Analysis of focal adhesion complexes and actin cytoskeleton**

Sterile glass coverslips were placed (Catalog : 100040N; Dutscher) into 24-well plates and coated with 350 μL fibronectin (7 μg/mL; Catalog : F1141, Merck) for 30 min at room temperature in a sterile hood. After removing the coating solution, coverslips were dried for 30 min. 4T1 WT and 4T1 LRP-1-knockout cell were seeded at $15.10^3$ cells per coverslip; HS578-T WT and HS578-T LRP-1-knockout cells were seeded at $5.10^4$ cells per coverslip. Cells were left to adhere for 24 h at 37 °C in a humidified 5% $CO_2$ incubator. Fixation was performed using a two-step paraformaldehyde (PFA) protocol to preserve morphology and reduce detachment. We added PFA directly to the culture medium to a final concentration of 2% and incubated for 5 min at room temperature, then washed once with PBS and completed fixation with 4% PFA for 10 min at room temperature. Coverslips were washed three times with PBS. Membrane permeabilization was achieved by incubating coverslips with 0.1% Triton X-100 in PBS for 4 min at room temperature. Non-specific binding sites were blocked with PBS containing 5% bovine serum albumin (BSA) for

30 min at room temperature. Coverslips were incubated overnight at 4 °C with anti-vinculin primary antibody diluted in PBS with 1% BSA. The next day, coverslips were washed five times with PBS (two quick washes followed by three washes of 5 min each). Secondary labelling was carried out at room temperature by incubating coverslips with Alexa Fluor 488 secondary antibody for 2 h together with Alexa Fluor 568-phalloidin for 1 h, both reagents were diluted in PBS containing 1% BSA. After staining, coverslips were washed five times with PBS (two quick washes and three washes of 5 min). Nuclei were counterstained with Hoechst dye according to the manufacturer's instructions and coverslides were washed twice with PBS before mounted on glass slides with an aqueous mounting medium. Observations were performed using an inverted microscope (EVOS FL Auto 2, Invitrogen, Thermo Fisher Scientific).

**Plasmin Activity Assay**

Conditioned media from cells cultured during 24 hours in phenol red-free and serum-free medium were analyzed for plasmin activity, using the chromogenic substrate S-2251™ (H-D-Val-Leu-Lys-pNA, Chromogenix, Diapharma, Westchester, USA) according to the manufacturer's instructions. For the assay, 20 µL of conditioned media were added to 145µL of Tris-HCl buffer (0.1 M, pH 7.8) and 20 µL of plasminogen (40 µg/mL) in a 96-well plate. The reaction was initiated by adding 20 µL of S-2251 substrate. Plates were incubated at 37°C in a spectrophotometer (Tecan), and absorbance at 405 nm was measured every 30 minutes for up to 18 hours.

**Protein extraction and Western blots:**

We performed Western blotting as previously described[13]. The antibodies used and their dilutions are listed in Additionnal_table_1. Band intensities were quantified by densitometry, normalized with the appropriate control , and are reported as mean ± SD. When comparing two groups, a two-tailed Student's t-test was used. Statistical significance is indicated in the text and figures by asterisks: $P < 0.05$ (*).

**Atomic force microscopy**

4T1 WT and LRP-1-knockout cells were seeded at $5.10^5$ cells per Wilco glass-bottom culture dish (Catalog : GWST-5040, WillCo Wells B.V., Amsterdam, The Netherlands). After plating, cells were left undisturbed for 24 h to allow proper adhesion and recovery before further experiments. Cells were imaged in PeakForce QNM™ mode on a Bioscope Catalyst™ (Bruker, Billerica, USA) with QNM-LC-CAL probes having a nominal spring constant of 0.1 N/m and a tip radius of 65 nm. We used the flowing setup: 2000 nm drive amplitude, 0.25 kHz drive frequency and scan rate of 0.3 to 0.5 Hz. Individual force curves were extracted and re-processed manually using a baseline correction algorithm with a contact-point based estimation to extract the Young's modulus. We spotted areas where the cell height is at least three times superior to the indentation depth, to avoid any influence of the substrate on the measurement. 10 force curves were captured per cell and the experiment was done in triplicate. In total, more than 170 curves were captured per condition (4T1 WT and KO-LRP-1). Prior to the experiment, the deflection sensitivity was estimated by extrapolating the linear portion of three force curves captured on a non-compliant sample in fluid and averaging the three values. The spring constant was given by the manufacturer but was double-checked by a thermal tune calculation achieved in the same medium (medium filter width: 3). Results were plotted as a violin plot. Statistical comparisons between groups were performed using the nonparametric Mann–Whitney test. Statistical significance is denoted as follows: $P < 0.0001$ (****).

**Measurements of Membrane Fluidity**

After being cultured in a Petri dish with RPMI media, the cells were incubated for 15 minutes at 37 °C after three rounds of washing with ice-cold PBS suspended in Laurdan solution (5 μM in PBS). A Zeiss (Oberkochen, Germany) LSM710 Meta confocal microscope with either a ×63 Plan Apochromat objective at a resolution of 132 nm.pixel−1, resulting in a small XY oversampling, was used to assess the fluorescence from confocal microscopy images of cells. 37 °C was the temperature that was set. Each detector has a BrightLine single-band bandpass filter in front of it, with the blue channel at 460/80 nm and the red channel at 540/50 nm. Photos were obtained using the Zen software program (ZEN 3.1 black edition LS.Ink).

GP was computed using the formula $GP = (I_{435} – I_{500})/( I_{435} + I_{500})$. Statistical comparisons between groups were performed using the nonparametric Mann–Whitney test. Statistical significance is denoted as follows: $P < 0.0001$ (****).

**Orthotopic TNBC mouse model**

All animal experimental procedures were conducted after obtaining authorization from the French Ministry of the Research (authorization #8140v7) and at an accredited scientific research animal facility (E-51-454-2). For orthotopic tumor implementation, eight weeks old BALB/c female mice (n=10 per group) (Janvier Lab, France) were injected with $5.10^5$ 4T1 WT or 4T1 KO-LRP-1 cells (100 µL of PBS) into mammary fat pad (MFP). Tumor growth was monitored every 3 days by measuring tumor length and width with calipers (tumor volume = ½ (length × $width^2$)), and mice were euthanized 26 days post-injection. Tumors were excised, measured, and either fixed in 4% paraformaldehyde (PFA) or processed for downstream analyses as described below. All experience are done on accordance with rules of local ethical committee (CEEA-51). Statistical comparisons between groups were performed using a two way ANOVA test. Statistical significance is denoted as follows: P<0,05 (*), P< 0,01 (**) and P < 0.0001 (****).

**Trichrome Masson Staining and Second Harmonic Generation imaging**

Second harmonic generation images were acquired using a Zeiss LSM710 microscope by using linear unmixing mode at 430 nm after excitation at 860 nm. Trichrome Masson staining was performed following the manufacturer's protocol (Diapath, 010210). Briefly, slides were deparaffinized, hydrated, covered with Gill's hematoxylin I for 5 minutes and saturated alcoholic picric acid for 5 minutes. Then, slides were rinsed with distilled water for 30 seconds and stained with Ponceau fuchsin for 10 minutes. Finally slides were treated with phosphomolybdic acid for 4 minutes and aniline blue for 2 minutes. Images were analyzed with ImageJ using color deconvolution tool plugin. The same threshold on the both groups of slides were applied and the results represent area covered by collagen on the total tumor area. Three different tumor for each groups were analyzed. Statistical comparisons between groups were performed using the nonparametric Mann–Whitney test. Statistical significance is denoted as follows: P < 0.05 (*).

**Cytokinome analysis**

The preparation of samples for cytokine profiling is described in fig_additional_4, same as secretomic (except that the concentration was carried out using Vivaspin with a cut-off of 3 kDa). Cytokine array was performed using Proteome Profiler Mouse XL (ARY028 R&D systems) and performed according to manufacturer's instructions and as in previous study[14]. Dot intensities were analysed using the QuickSpots software and following manufacturer's instructions. Only the cytokines showing the most significant and robust changes, as determined by strict exclusion criteria from two independent experience with excluding weak or inconsistent spots, were included in the figures presented.

### Additionnal_table_1

#### Antibody list

| Antibodies | Reference | Methods | Dilution |
|---|---|---|---|
| LRP-1 | HPA004182 | Immunohistochemistry | 1:500 |
| HRP/DAB IHC detection kit | ab236466 | Immunohistochemistry | Ready to use |
| NCAM1 | ab237708 | Immunohistochemistry | 1:1000 |
| CD8 | ab316778 | Immunohistochemistry | 1:500 |
| CD4 | ab183685 | Immunohistochemistry | 1:150 |
| CD3 | ab135372 | Immunohistochemistry | 1:100 |
| LRP-1 | Merck, HPA004182 | Immunohistochemistry on spheroids | 1:100 |
| E-cadherin | 24E10 | Immunohistochemistry on spheroids | 1:800 |
| N-cadherin | D4R1H | Immunohistochemistry on spheroids | 1:125 |
| Ki-67 | ab16667 | Immunohistochemistry on spheroids | 1:200 |
| LRP-1 | EPR3724 | Western-blot | 1:10 000 |
| β-actin | sc-47778 | Western-blot | 1:1000 |
| LOXL4 | sc-365822 | Western-blot | 1:200 |
| MMP14 | NBP2-67415 | Western-blot | 1:1000 |
| Goat anti-Rabbit IgG (H+L) Secondary Antibody, DyLight™ 800 4X PEG | SA5-35571 | Western-blot | 1:10 000 |
| Goat anti-Mouse IgG (H+L) Secondary Antibody, DyLight™ 800 4X PEG | SA5-35521 | Western-blot | 1:10 000 |
| Goat anti-Mouse IgG (H+L) Secondary Antibody, DyLight™ 680 - | 35518 | Western-blot | 1:10 000 |
| Anti-rabbit IgG, HRP-linked Antibody | 7074 | Western-blot | 1:10 000 |
| Anti-mouse IgG, HRP-linked Antibody | 7076 | Western-blot | 1:10 000 |
| vinculin | 700062 | Immunofluorescence | 1:100 |
| Alexa Fluor 488 secondary antibody | A-21206 | Immunofluorescence | 1:1000 |
| GR1 APC | 108455 | flow cytometry | 1:100 |
| CMHII PE | 107608 | flow cytometry | 1:100 |
| PD-1 BV421 | 135217 | flow cytometry | 1:100 |
| CD44 BV605 | 103047 | flow cytometry | 1:100 |
| Ly6C PercpCy5.5 | 128011 | flow cytometry | 1:100 |
| CD11b PE/Cy5 | 101210 | flow cytometry | 1:100 |
| CD8 BV785 | 100749 | flow cytometry | 1:100 |

| | | | |
|---|---|---|---|
| Tim3 BV711 | 119727 | flow cytometry | 1:100 |
| F4/80 Pe/Cy7 | 123113 | flow cytometry | 1:100 |
| CD45 BV510 | 103137 | flow cytometry | 1:100 |
| CD4 AF700 | 100536 | flow cytometry | 1:100 |
| NKp46 BV711 | 137621 | flow cytometry | 1:100 |
| TCRβ PercpCy5.5 | 560657 | flow cytometry | 1:100 |

#### **staining list**

| Staining | Reference | Methods | Dilution |
|---|---|---|---|
| Alexa Fluor 568-phalloidin | A12380 | Marquage cytosquellette d'actine | 1:40 |
| Zombie Green | 423111 | flow cytometry | 1:100 |
| Hoescht | 62249 | Immunofluorescence | 1:1000 |
| Gill's hematoxylin 2 | 05-0614/L | HES staining | Ready to use |
| 0.5% alcoholic eosin G | 05-10009/L | | Ready to use |
| saffron | F/SAFRAN | | 0.5% |
| Trichrome Masson stain Kit | 010210 | Staining of connective tissue, collagen, reticular and muscle fibers | Ready to use |
